# Disentangling the relative roles of vertical transmission, subsequent colonizations and diet on cockroach microbiome assembly

**DOI:** 10.1101/2020.07.01.183558

**Authors:** Kristjan Germer, Justinn Renelies-Hamilton, David Sillam-Dussès, Kasun H. Bodawatta, Michael Poulsen

## Abstract

A multitude of factors affect the assemblies of complex microbial communities associated with animal hosts, with implications for community flexibility, resilience and long-term stability; however, their relative effects have rarely been deduced. Here, we use a tractable lab model to quantify the relative and combined effects of parental transmission (egg case microbiome present/reduced), gut inocula (cockroach vs. termite gut provisioned), and varying diets (matched with gut inoculum source) on gut microbiota structure of hatchlings of the omnivorous cockroach *Shelfordella lateralis* using 16S rDNA amplicon sequencing. We show that the presence of a pre-existing bacterial community via vertical transmission of microbes on egg cases reduces subsequent microbial invasion, suggesting priority effects that allow initial colonizers to take a stronghold and which stabilize the microbiome. However, the subsequent inoculation sources more strongly affect ultimate community composition, with distinct host-taxon-of-origin effects on which bacteria establish. While this is so, communities respond flexibly to specific diets that consequently strongly impact community functions predicted using PICRUSt2. In conclusion, our findings suggest that inoculations drive communities towards different stable states depending on colonization and extinction events, through ecological host-microbe relations and interactions with other gut bacteria, while diet in parallel shapes the functional capabilities of these microbiomes. These effects may lead to consistent microbial communities that maximize the extended phenotype that the microbiota provides the host, particularly if microbes spend most of their lives in host-associated environments.

**Contribution to the field:** When host fitness is dependent on gut microbiota, microbial community flexibility and reproducibility enhance host fitness by allowing fine-tuned environmental tracking and sufficient stability for host traits to evolve. Our findings lend support to the importance of vertically transmitted early-life microbiota as stabilizers through interactions with potential colonizers that may contribute to ensuring that the microbiota aligns within host fitness-enhancing parameters. Subsequent colonizations are driven by microbial composition of the sources available, and we confirm that host-taxon-of-origin affects stable subsequent communities, while communities at the same time retain sufficient flexibility to shift in response to available diets. Microbiome structure is thus the result of the relative impact and combined effects of inocula and fluctuations driven by environment-specific microbial sources and digestive needs. These affect short-term community structure on an ecological time scale, but could ultimately shape host species specificities in microbiomes across evolutionary time, if environmental conditions prevail.

## Introduction

Intricate associations between animal hosts and their gut microbiota are vital for the evolution and persistence of many animal hosts (1, 2). These microbial symbionts facilitate a multitude of functions associated with host nutrient management, immunity and development (1, 3), and ultimately impact host adaptation and diversification across environments and dietary niches (e.g., 4-8). When hosts traits select for specific microbial functions, these can be considered the extended phenotype of the host (9–12). Selection should optimally involve getting a microbiota that is both flexible (i.e., containing environment-specific strains that are likely to enable degradation of environment-specific nutrients and toxins) and consistent (i.e., similar under a defined set of circumstances) rather than subject to random fluctuations (3, 13–15).

The composition of complex gut microbial communities in many mammals, birds, and insects (16) are driven by host taxonomy (17–20), diet (4, 5), vertical (parent-offspring) transmission (21), and environmental inputs (22), including transmission from conspecifics through social behaviors (23) [e.g., coprophagy (24, 25) and trophallaxis (24)]. Early-life microbial colonizations, including vertical transmission and environmental inputs, will have a disproportionate effect on the microbiota (priority effects); with subsequent positive (facilitation) and negative (competition) interactions between community members affecting ultimate composition (26–29). In addition, gut physiology and diet impose strong filters that limit what microbes can establish and ultimately diversify with host species (28, 30, 31). Diets will on average be more similar between individuals of the same host species than between species, and they may hence contribute to microbiota consistency within species on an ecological time scale (32) and ultimately long-term association across evolutionary time (2). While studies exploring the impact of host phylogeny (e.g., 33), diet (33, 34), or microbial inocula (31) on microbiota structure are many, studies that allow testing of their relative and combined effects are few.

To contribute to closing this knowledge gap, we quantify the relative and combined effects of transmission (with or without disrupted vertical transmission), environmental microbial sources, varying diets and host specificity of the above on gut microbiota structure in hatchlings of the omnivorous cockroach *Shelfordella lateralis* (Turkestan cockroach; Blattodea: Blattidae). Cockroaches are excellent models for testing the impact of these factors, due to their abundant, diverse and consistent yet dynamic microbiota, which is amenable to antimicrobial and dietary manipulations (34–37). We exposed developing nymphs with or without access to bacterial communities on the ootheca (the egg cases they emerge from, a potential source of parental gut microbes), as whether the importance of this indirect vertical transmission route is similar to other cockroaches remains unknown (38, 39). Next, we expose both groups of nymphs to con- or allospecific microbial inocula (cockroach vs. termite origins) and the corresponding diets of the two species (omnivorous vs. specialized fungus). We chose these species, because termites are social cockroaches and fungus-growing termites in particular share many cockroach gut bacterial lineages (17, 40). In doing so, we show that ecological interactions are important for microbial assemblage and that removing the initial microbiome, even if later reinoculated, will strongly affect microbiome consistency even if most microbes are shared. Furthermore, subsequent differences in available microbial inoculum sources and diets in combination drive ultimate microbiota structure and function.

## Results

### Antimicrobial treatment effectively reduces host microbiome

We confirmed that antimicrobial treatment with peracetic acid of ootheca reduced bacterial diversity in the initial microbiota of developing *S. lateralis* nymphs, using an established protocol that does not appear to negatively affect the cockroaches (31, 41). First, we compared the number of bacterial Colony Forming Units (CFUs; Fig. S1) that emerged on growth medium after plating of cockroaches from treated/untreated ootheca and found that the number was significantly lower in treated individuals (mean CFUs/sample ± SE: 5.08 ± 2.48 and 55.33 ± 20.34, respectively; WelchADF; WJ = 5.781, df = 1, p = 0.0434; Fig. S1). Secondly, we performed MiSeq amplicon sequencing of one cockroach per treated or untreated ootheca used in the main experiment (see below) and found a reduction in Amplicon Sequence Variant (ASV) diversity by 36.8% in nymphs from treated ootheca (Kruskal-Wallis; χ^2^ = 4.333, df = 1, p = 0.0374; Table S3), but no effect of treatment on community richness (χ^2^ = 2.077, df = 1, p = 0.1495) or beta diversity (PERMANOVA_10,000 permutations_; F_1_ = 1.268, p = 0.2873) (Table S3). As expected, the obligate endosymbiont *Blattabacterium* was unaffected by treatment (cf., 42), which was interestingly also the case for an *Alislipes*_III ASV (Fig. S3; Table S3). All other ASVs were reduced in relative abundance, often below the level of detection (Fig. S3). In summary, although treatment with peracetic acid did not sterilize developing nymphs, microbiome diversity was substantially depleted.

### The combined effects of antimicrobial treatment, inoculum and diet on microbiome structure

Using 16S rDNA MiSeq amplicon sequencing of whole guts after a 28-day treatment period, we tested for the combined effects of the depletion of the microbiome on the ootheca (presence/absence of peracetic acid treatment), environmental inocula (*S. lateralis* vs. *Macrotermes subhyalinus* gut provided on the first day of the experiment), genetic background (ootheca identity), and diet (omnivorous dog food vs. specialized fungus diet provided *ad libitum)* (see Fig. 1 for the experimental setup, and Materials and Methods for details). Ordination analyses (Fig. 2) revealed grouping in NMDS space that were consistent with strong effects of inoculum (PERMANOVA_10,000 permutations_; F_1_ = 15.24, p_adj_ < 0.0001, R^2^ = 0.1145), diet (F_1_ = 8.734, p_adj_ < 0.0001, R^2^ = 0.0657), and minor effects of antimicrobial treatment (F_1_ = 3.422, p_adj_ < 0.0001, R^2^ = 0.0257) and host genetic background (ootheca origin) (F_10_ = 1.561, p_adj_ < 0.0001, R^2^ = 0.1174). Given the absence of clustering in the NMDS plot (Fig 2B), the variation explained by the latter is likely due to the number of groups this factor contained and it should thus be interpreted cautiously (Figs 2B; S4; Table S5).

**Figure 1.**
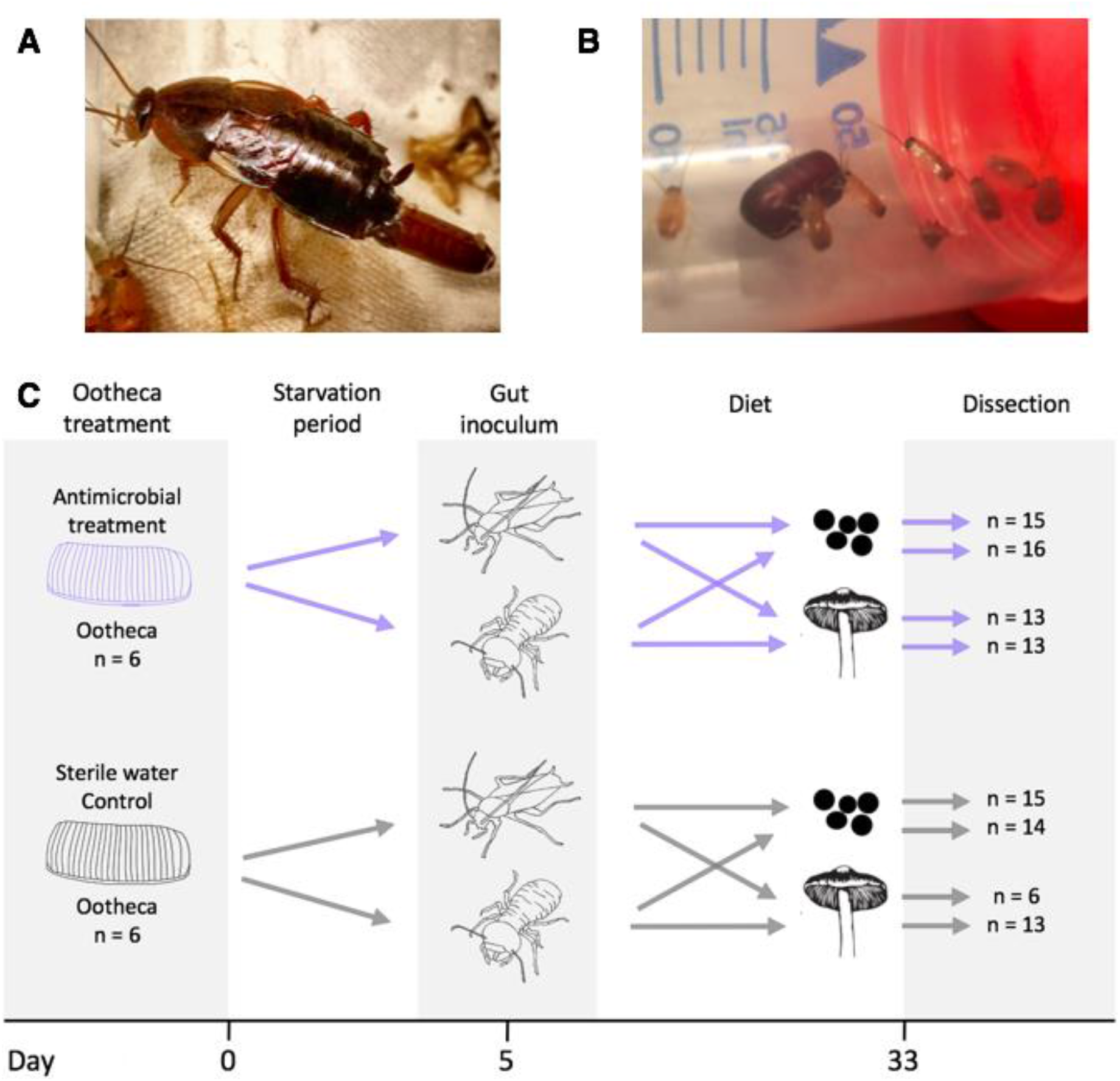
A. *Shelfordella lateralis* laying an ootheca with eggs. B. Newly hatched *S. lateralis* nymphs with their ootheca (Photos KG). C. Schematic of the main experimental setup and timeline; silhouette of termites and mushrooms by Rafael R. da Costa, cockroach and dog food by KHB. Cockroaches hatched on day 0 and a single cockroach was removed to assess antimicrobial treatment effectiveness on day 1. Blue arrows represent the ootheca treated with peracetic acid to deplete the microbiome and grey arrows represent controls. Sample sizes are indicated with “n”.

**Figure 2.**
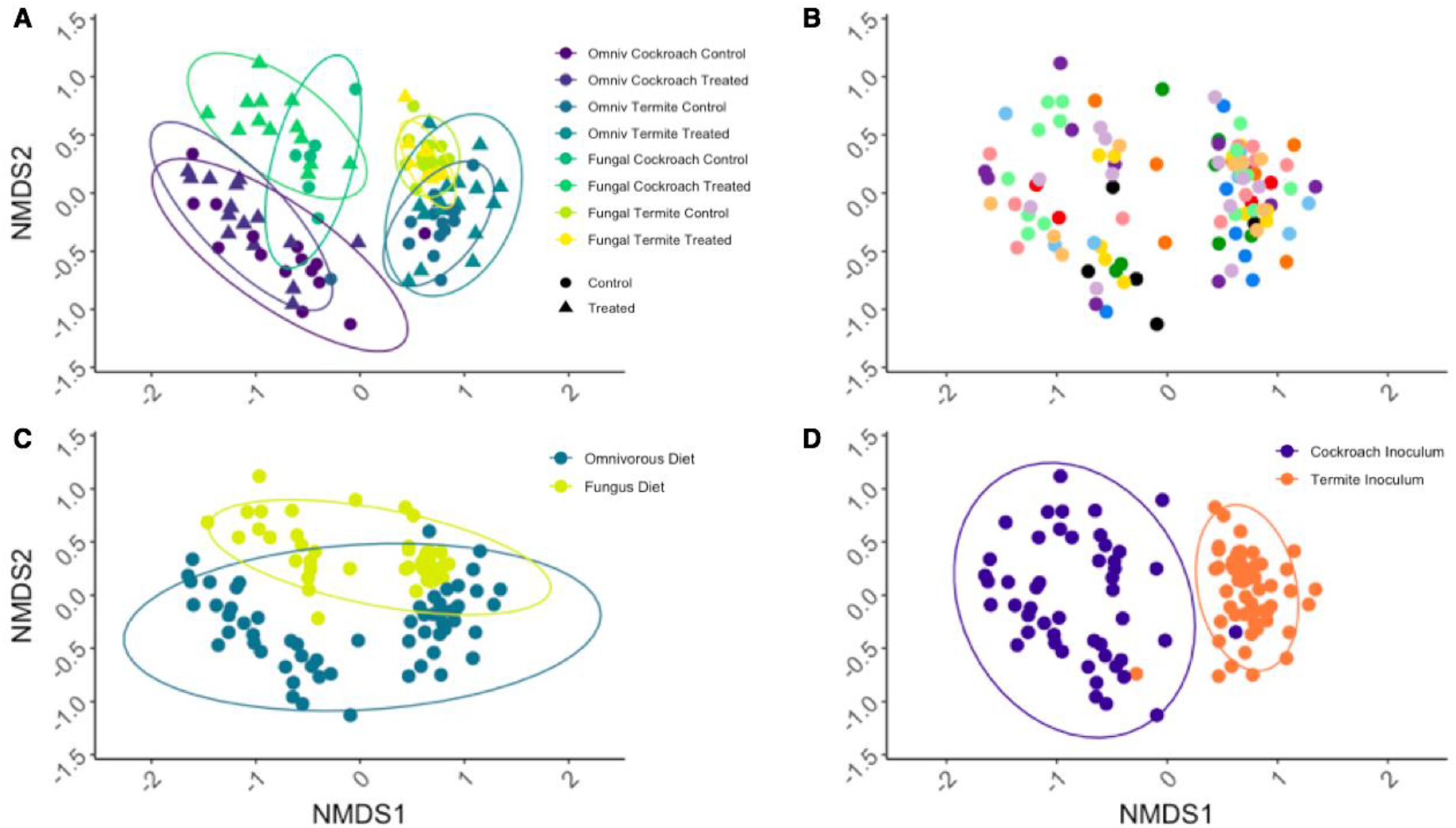
Bray-Curtis distance NMDS ordination plot with identical coordinates in all panels, stress = 18.2%. In A, triangles indicate peracetic acid-treated cockroaches and circles controls. Colour indicates treatment groups: cockroach gut and omnivorous diet in purple, termite guts and omnivorous diet in dark green, cockroach guts and fungus diet in teal, and termite guts and fungus diet in yellow (see Fig. 3). Panel B is colored by cockroach genetic background, panel C by diet: omnivorous in blue and fungal in ochre, and panel D by microbial inoculum: cockroach gut in purple and termite gut in orange. Ellipses indicate 95% confidence intervals.

Gut microbiome alpha diversity was affected by diet, inoculum, and antimicrobial treatment, but unaffected by cockroach genetic background (Fig. 3A; S5; Table S4). Antimicrobial-treated cockroaches regain a more diverse gut microbiome following microbial inoculation, especially when inoculum is sourced from conspecific hosts (ANOVA; F_1_ = 6.522, p = 0.0122; Fig. 3A). Cockroach inocula produced, on average, a 69.7% more diverse community than termite inocula, indicating that conspecific inoculum allows more microbial gut colonization than allospecific ones (F_1_ = 17.12, p = 8e-5). Conversely, however, the fungal diet sustained, on average, a 17.7% more diverse community than the omnivorous diet the cockroaches normally feed on (F_1_ = 8.547, p = 0.0044). In line with the beta diversity results, inoculum had the largest effect size (0.4701), followed by diet (0.3366) and antimicrobial treatment (0.2918) (Table S4).

**Figure 3.**
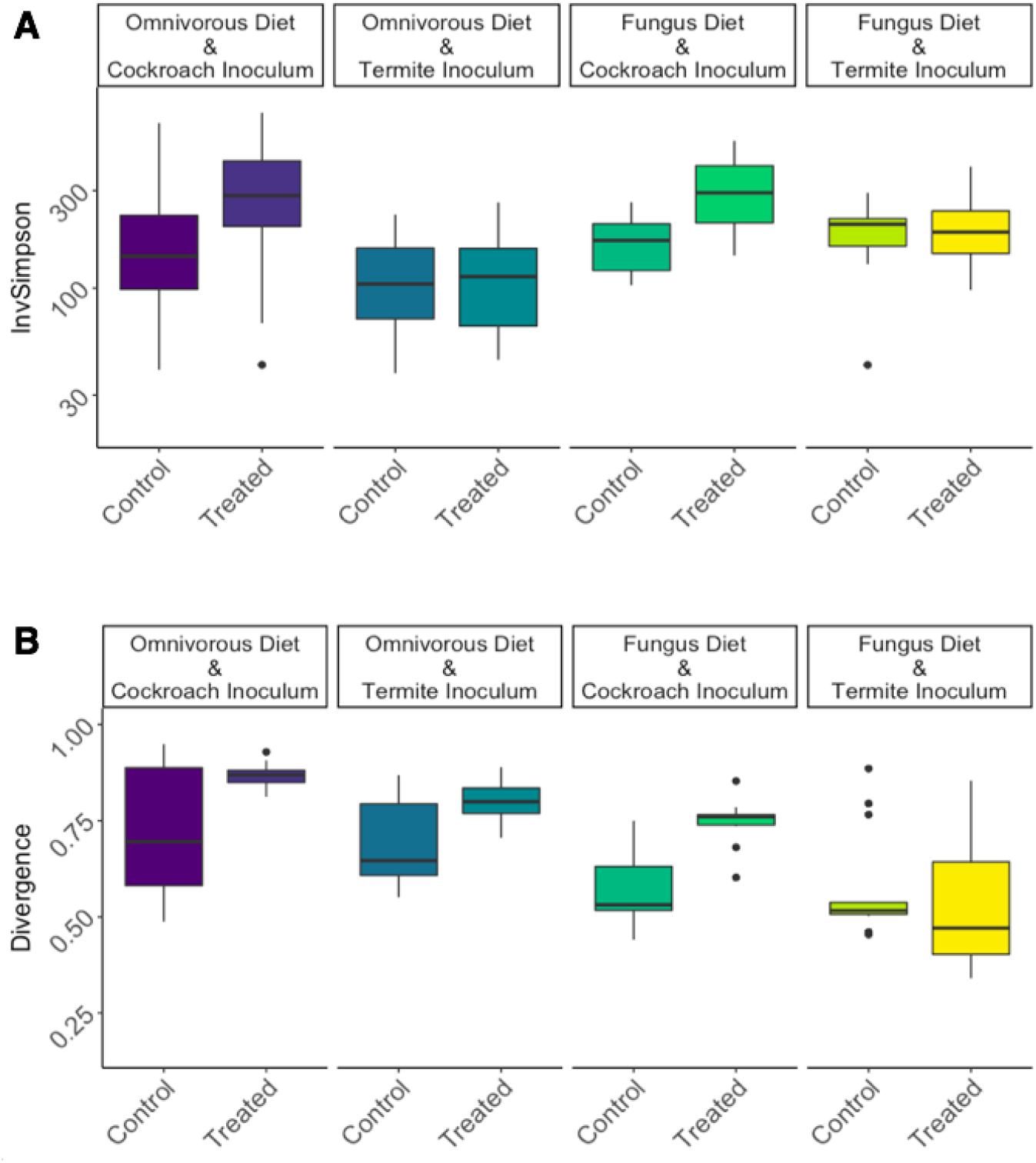
(A) Inverse Simpson diversity index for each treatment group on a log10 scale. (B) Microbiome divergence (inverse metric for consistency) calculated as beta diversity between each sample in a group and the representative median abundance of each microbe in that group. Horizontal lines indicate medians, hinges indicate first and third quantiles. Colouration of treatment groups corresponds to that of Fig. 2A.

To explore whether disrupting the early-life microbiome and varying microbial inocula and diets affected how consistent microbiomes were within treatments, we modelled between sample-divergence within groups (see Materials and Methods; 43). Antimicrobial treatment produced a sharp decrease in microbiome consistency (WelchADF; WJ = 16.81, df = 1, p = 0.0002; Fig. 3B; Table S6), while inoculation with cockroach guts decreased microbiome consistency compared to termite guts (WJ = 8.905, df = 1, p = 0. 0046) as did an omnivorous diet compared to a fungal diet (WJ = 45.075, df = 1, p = 2.672e-8). Lastly, there was a significant compensatory or negative interaction between inoculum and antimicrobial treatment. Antimicrobial-treated cockroaches fed on cockroach inocula were significantly more variable than controls, and antimicrobial-treated individuals inoculated with a cockroach microbiome experienced increased divergence than individuals inoculated with termite microbes (WJ = 7.048, df = 1, p = 0.0109).

### Taxa effects driving the community-level differences associated with treatment

To determine what taxa were affected most by our treatments, we used an ANOVA-like multivariate model (44, 45) while accounting for the independent effects of antimicrobial treatment, diet, and inoculum. Twenty genera increased significantly in relative abundance in groups receiving a cockroach inoculum, five with a termite inoculum, and seven in cockroaches on a fungus diet (Fig. 4; Table S7). Interestingly, no taxa increased significantly in relative abundance in cockroaches fed the omnivorous diet. A set of taxa increased in abundance with the cockroach inoculum: Porphyromonadaceae_Cluster_V termite_cockroach_cluster, *Alistipes_*II, III and IV, Ruminococcaceae insect_cluster and termite_cockroach_cluster, *Mucispirillum*, Veillonellaceae uncultured_7, Lachnospiraceae gut_cluster_13, Desulfovibrionaceae Gut_cluster_II and Synergistaceae termite_cockroach_cluster (Fig. 4). All of these taxa have diversified within the Blattodea to some extent and are found in both fungus-growing termites and cockroaches (30, 46).

**Figure 4.**
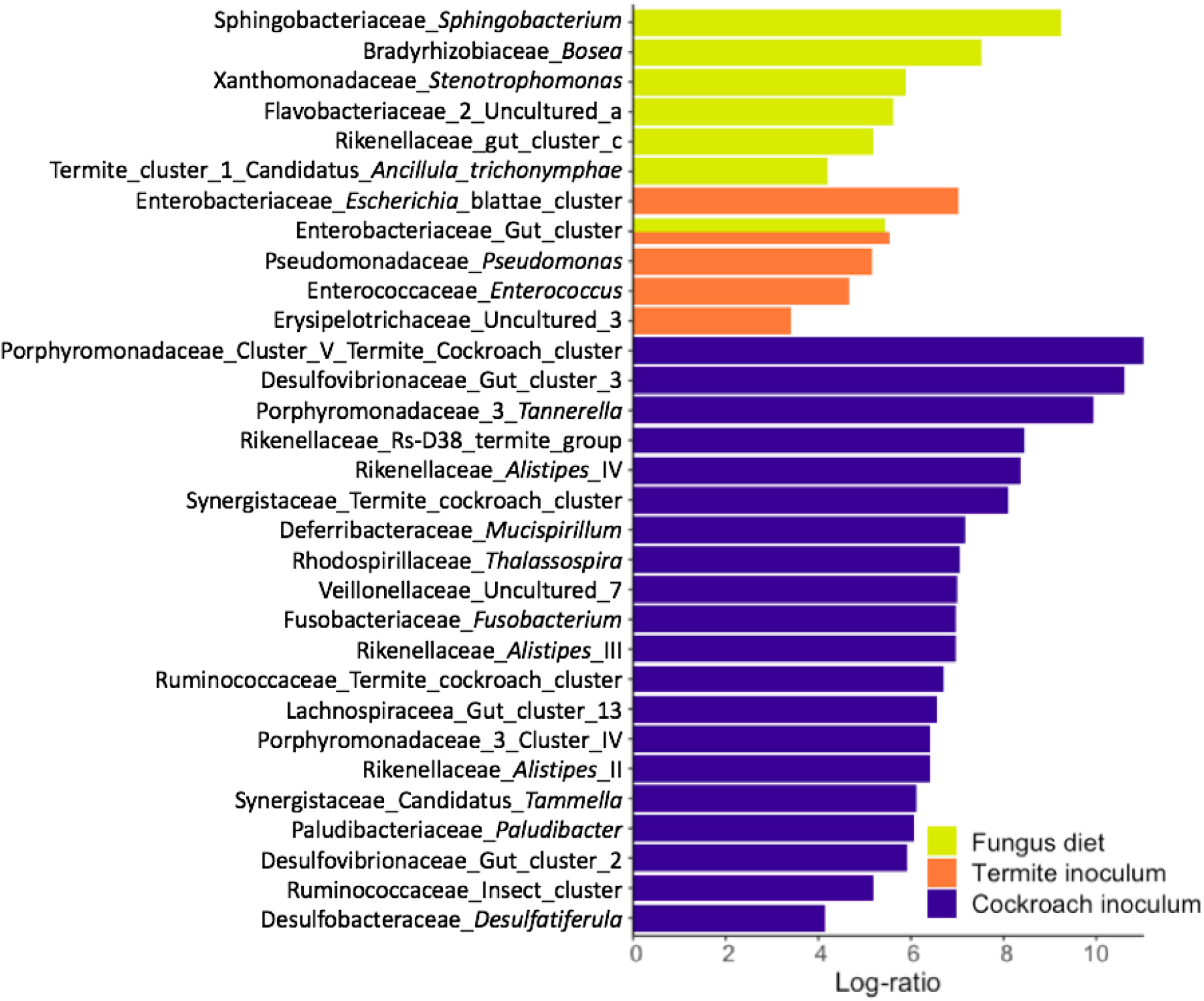
Log-ratio (effect size) increase in genera significantly correlating with inoculum or diet compared to their alternative. Calculated with a multivariate Aldex2 generalized linear mixed model. All plotted taxa have Bonferroni-Hochberg adjusted p values below 1e-5.

### Functional prediction changes associated with changing inoculum and diets

To discern any changes in predicted functional capabilities of bacterial communities as a whole, we employed PICRUSt2 on the 16S rRNA sequences (47) and subsequently identified predicted microbial metabolic pathways using the MetaCyc database (48). Overall, there were 406 predicted pathways in the cockroach gut microbiomes. Although microbial function was more conserved than microbiome composition (Figs. 5; S7; Table S9), consistent with previous studies (e.g., 49), there were clear differences in response to our treatments (Table S10). Diet had a strong and significant impact on the metabolic profiles, explaining 22.2% of the observed variation (PERMANOVA_10,000 permutations_; F_1_ = 37.08, p_adj_ = 1e-5) (Fig. 5). Gut inoculum (F_1_ = 10.30, p_adj_ =5e-5), antimicrobial treatment (F_1_ = 3.081, p_adj_ = 0.0185), and genetic background (F_10_ = 2.631, p_adj_ = 0.0002) also had significant impacts, yet the variation explained by these factors were lower (only 6.2, 1.9 and 15.8%, respectively) (Fig. 5).

**Figure 5.**
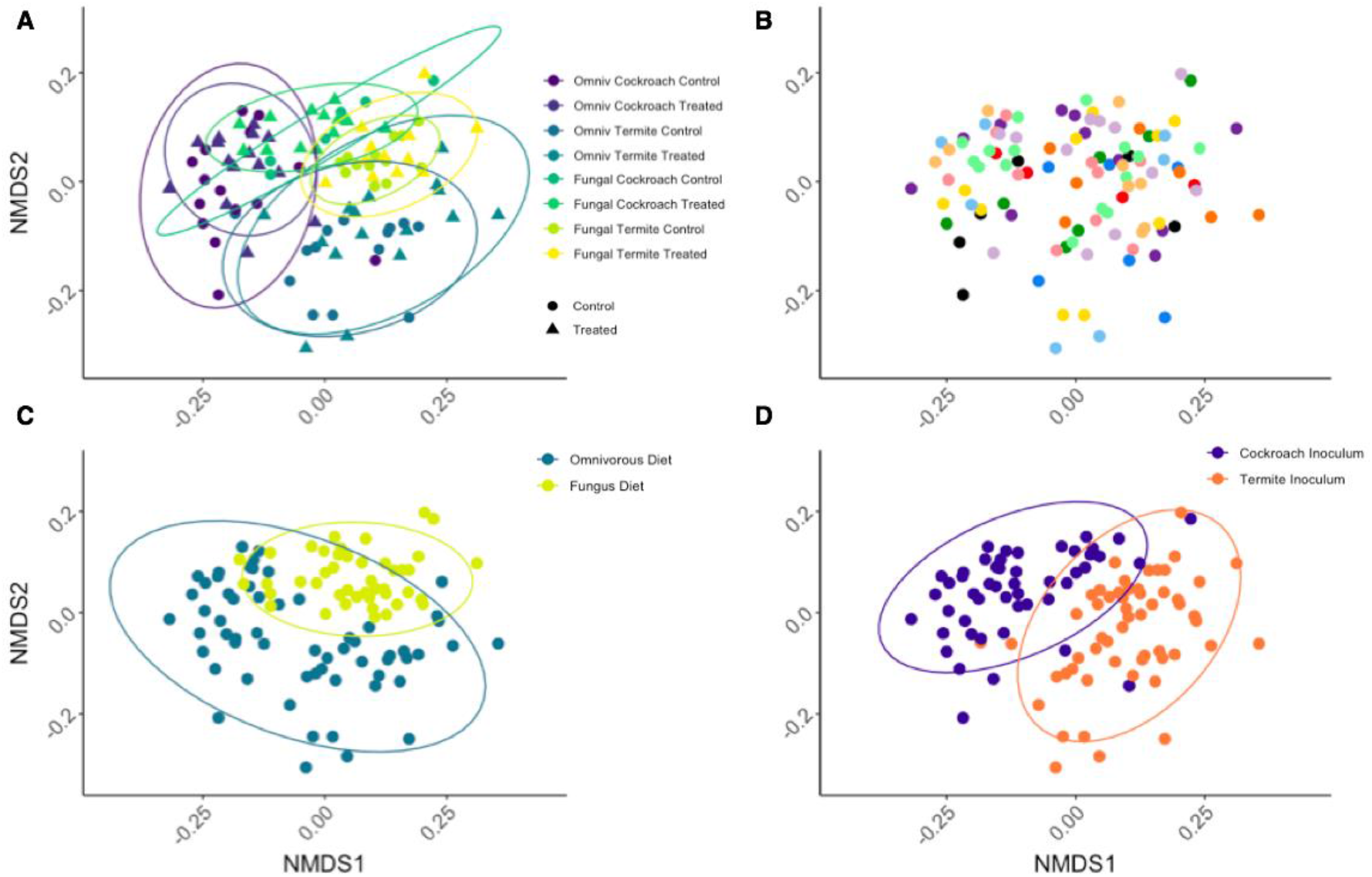
NMDS plot of the predicted bacterial metabolic pathways (PICRUSt2) of gut microbiomes of cockroaches with different gut inocula and diets. A: all groups; B: genetic background; C: diet; D: inoculum; coloration of treatment groups as in Fig. 2. Ellipses indicate 95% confidence intervals.

The microbiome responded to a fungal diet by increasing slightly MetaCyc pathway richness and diversity (WelchADF; WJ = 192.8, df = 1, p < 1e-10; WJ = 24.95, df = 1, p = 6.666e-6; Figs. S8A; S8B; Tables S11; S12). Termite inoculum increased pathway diversity, but not richness (WJ = 9.468, df = 1, p = 0.0033) while antimicrobial treatment increased pathway richness in hosts inoculated with cockroach guts (WJ statistic = 5.115, df = 1, p = 0.0268). Subsequently, we analyzed functional divergence as we did for microbiome composition divergence (see Materials and Methods and above). Mirroring composition divergence, the omnivorous diet produced a more variable functional microbiome than a fungal diet (WJ = 12.42, df = 1, p = 0.0008; Fig. S8C; Table S13). Interestingly, an interaction between diet and inoculum indicated that when these are matched (i.e., cockroach inoculum together with omnivorous diet, and termite inoculum with fungal diet), there is significantly lower functional divergence than when they are mismatched (WJ = 8.251, df = 1, p = 0.0056).

To test for differences in predicted pathway abundances between treatments, we ran Aldex2 analyses (44, 45) (Table S14 gives the full results, Fig. S9 all significant pathways and Fig. 6 significant pathways that increased more than four-fold in centered-log ratio abundance). A total of 266 pathways were significantly enriched across treatment groups, with 26 being in cockroaches on a fungus diet, 138 on the omnivorous diet, 70 with termite inoculum, and 57 with cockroach inoculum. Of these, only 30 overlapped between groups: 26 between cockroach inocula and omnivorous diet, one between cockroach inocula and fungal diet, one between termite inocula and omnivorous diet, one between termite inocula and fungal diet, and one between antimicrobial treated cockroaches and the omnivorous diet (Table S14). Among pathways increasing in cockroaches on a fungal diet, 24 were more than 4-fold increased, with the most increased pathway being chitin breakdown, inevitably due to the abundance of this carbon source in the diet. Pathways increasing in abundance also included several amino acid degradation pathways (three out of four were for L-tryptophan), as well as a series of pathways for aromatic compound degradation (Fig. 6). The role of termite gut bacteria in the breakdown of lignin-derived aromatic compound (50) likely caused seven pathways for aromatic compound degradation to increase more than 4-fold when cockroaches were offered a termite gut inoculum. Three pathways associated with proteinogenic amino acid degradation were also increased in this group. Predicted proteinogenic amino acid degradation also increased in cockroaches fed a cockroach inoculum, as did several pathways for sugar biosynthesis were predicted. Interestingly, the omnivorous diet did not lead to increases of any pathways more than 4-fold, although a high number of pathways did significantly increase in cockroaches on this diet (Table S14).

**Figure 6.**
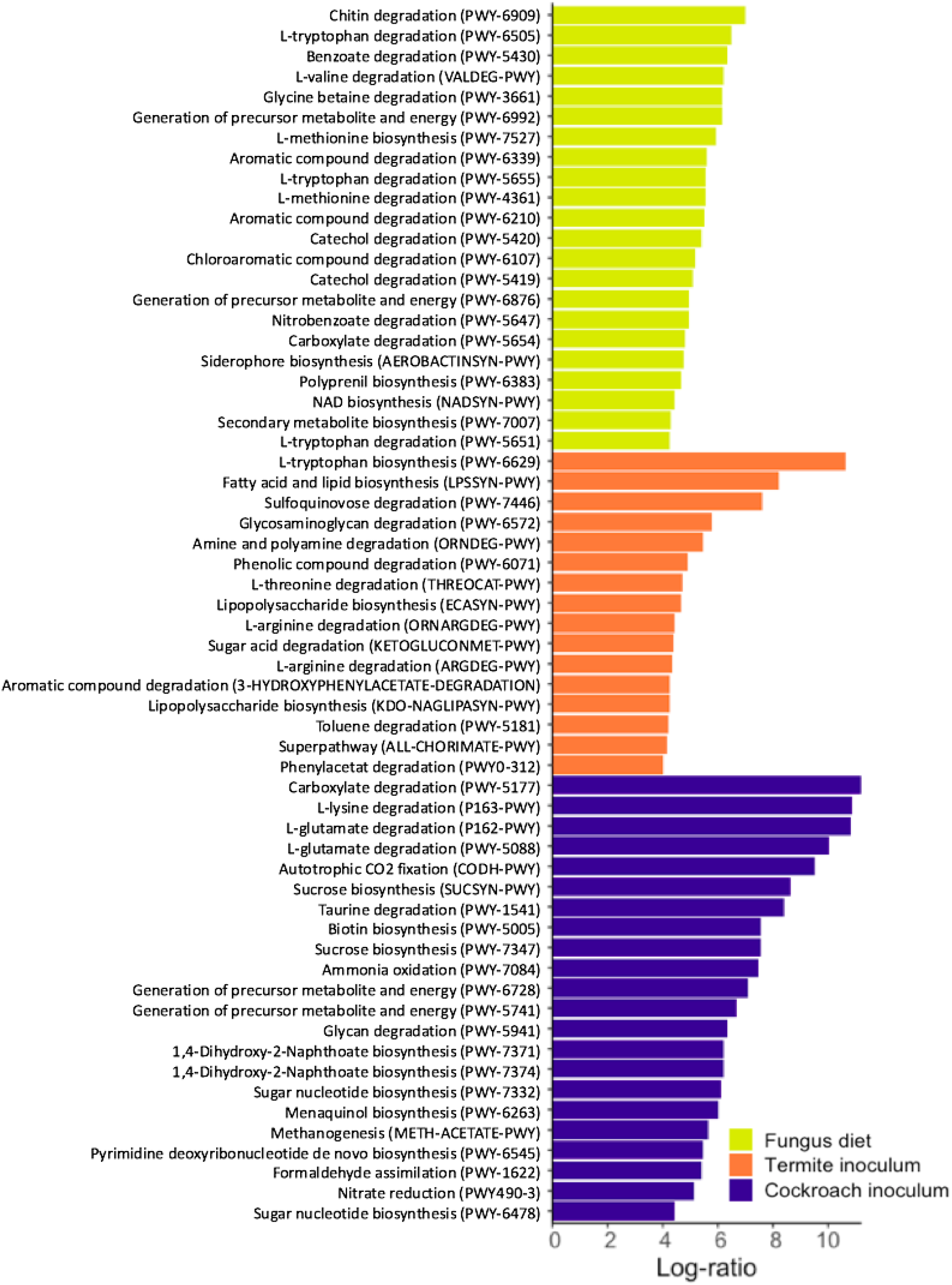
Log-ratio (effect size) increase in functional metabolic categories (pathways in brackets) that significantly correlated with inoculum or diet compared to their alternatives, derived from a multivariate Aldex2 generalized linear mixed model. Only log-ratios above four are shown, see Fig. S9 for all significant pathways and Table S14 for the full results.

Finally, we tested whether any treatment enriched broader functional categories in differentially abundant MetaCyc pathways (Table S15). Overall, the omnivorous diet treatment led to enrichment of biosynthesis pathways (Fisher’s exact test, p_adj_ = 3.748e-9), particularly the major categories of amino acid (p_ad_j = 1.945e-8) and nucleoside/nucleotide (p_adj_ = 2.276e-8) biosynthesis, and the functional groups of proteinogenic amino acid (p_adj_ = 1.806e-5), and purine nucleotide biosynthesis (p_adj_ = 0.0436) and degradation (p_adj_ = 0.0002). In cockroaches on a fungal diet, however, degradation, utilization, and assimilation pathways were enriched (p_adj_ = 0.0449). The termite inoculum enriched phenolic compound degradation (p_adj_ = 0.0493) and fatty acid (p_adj_ = 0.0456) and quinol/quinone (p_adj_ = 0.0017) biosynthesis.

## Discussion

*Shelfordella lateralis* cockroach hatchlings emerging from their ootheca rely on independently acquiring all bacterial inocula to colonize their guts, except for vertically-transmitted *Blattabacterium* endosymbionts (cf., 42). Our findings suggest that the first microbial colonization is through bacteria present on the ootheca, implying an extent of vertical transmission. This impacts subsequent invasions by microbes, consistent with how early-life microbiota has a disproportionate impact on community assembly; in part due to priority effects allowing initial colonizers to take a stronghold (28, 29, 51, 52). We found this priority effect to only slightly affect which microbiome the cockroaches had a month later when the experiment was terminated. However, oothecal microbes in the gut did compete with subsequent colonizers; thus, guiding microbiome assembly towards a simpler community compared to when ootheca microbes were depleted (Fig. 3A). Each colonizing microbe can stochastically lead community assembly in diverging directions and towards different stable states (cf., 52). The more stochastic this process is, the noisier the community assembly is expected to be. Competition offered by the very-early-life inoculum, and the consequent decrease in colonizing microbes, allows community assembly to be more reproducible (cf., the consistency patterns in Fig. 3B) as has been elegantly shown before in the simpler communities hosted by *Drosophila* flies (52). This points to a stabilizing role of semi-vertically transmitted taxa in *S. lateralis,* and, consequently, to the contribution of host traits enhancing initial inoculum towards ensuring that the microbiota develops reproducibly. This interpretation is in line with previous work elucidating the importance of competition for microbiome stability (53, 54), an effect which seems to start at the very earliest stages of microbial assembly in both simple (52) and complex (this study) host-associated microbial communities.

After 28 days of treatment, cockroach microbiomes shifted most prominently according to the single event of environmental microbial inoculum and, secondarily, according to the continuously-provisioned diet. This led to four distinct clusters in NMDS space that correspond to the combination of diet and inoculum (Fig. 2A). The strong impact of inoculum implies that a single experimental introduction event with microbes able to colonize a gut system can lead to remarkably distinct microbiome structures. This underlines the impact the gregarious lifestyle of *S. lateralis* combined with coprophagous behavior (24) has by offering ample opportunities for microbial uptake from conspecifics. This is consistent with recent work showing that fecal pellets in the diet can restore microbiomes after antibiotic treatment of *Blatta germanica* cockroaches (38). Microbe-sharing benefits of group-living (24, 55–58) can thus reliably assure horizontal transfer of gut microbes from the environment and eliminate the need for vertical transmission for the acquisition of a taxon-specific microbiota. Such behavioral microbiota filtering can contribute to patterns of phylosymbiosis, i.e., matches between host phylogeny and microbial community composition (2, 58), and help explain co-divergence between Blattodea and some of their microbial symbionts (59).

Host-symbiont adaptations are expected to allow conspecific inocula to provoke higher colonization rates than allospecific inocula, and this is what we observe. Cockroach inocula have more strains that are able to colonize guts than a related termite microbiome, despite hosting a markedly less diverse gut microbiome than the termite species (17). Consistent with this assertion, taxa that have diversified within Blattodea, and whose respective strains are shared by cockroaches and termites (30, 46, 59), display improved colonization success and increase in relative abundance when originating from a cockroach compared to a termite gut (Fig. 4), in support of host-symbiont adaptations (cf., 31). However, this increase in colonization rate by conspecific symbionts may lead to a decrease in microbiome consistency (Fig. 3B). Since more taxa are able to establish, and colonization is stochastic, a conspecific inoculum leads community assembly in more divergent directions and towards different stable states (cf., 52). The stabilizing role of semi-vertically transmitted bacteria is consequently also most prominent under a conspecific diet and inoculum, and it is absent with mismatched diet and inoculum (Fig. 3B). This further emphasizes the importance of host-microbe association specificity, in so far as their effects on ecosystem stability take place only while interacting with a host-specific community or under the influence of the diet hosts have adapted to.

While microbial inoculum had the strongest impact on community structure, functional inference tells a different story: diet had by far the largest impact on predicted microbial metabolism, explaining over three-fold more variation than inoculum (Fig. 5, Table S10). The predicted ability of the microbiome to digest diet compounds generally shifted from nitrogen-rich nutrients on the omnivorous diet to fatty acid and aromatic compounds on the fungal diet. This could be simply reflecting nutrients in diets, or diet could select for taxa able to degrade compounds which are not found in the diets we provide, but that are usually associated with such diets. Several taxa increased in abundance in response to a fungus diet, and are known to degrade fungus-derived polysaccharides in fungus-growing termites (e.g., Enterobacteriaceae, *Stenotrophomonas* and *Pseudomonas;* 60, 63; Fig. 4). The fact that diet enriched such taxa and functional groups irrespective of which inoculum they received (Figs. 5, 6) suggests substantial redundancy in cockroach and termite microbiome functions, a well-established phenomenon across even disparate microbiomes (49). Therefore, inoculum and diet play their roles at different levels: colonizations are largely driven by what microbes are present in the environment and their ability to establish within hosts, while microbiome function is largely driven by what diet this host receives.

Although PICRUSt2 analysis should be evaluated with caution as it is extrapolative from 16S rRNA sequences, we find plausible specific functional changes that are relevant for specific diets, such as chitin degradation in cockroaches on the fungal diet and differences in amino acid metabolism across diets (Figs. 5, 6). Other predicted enrichments are more likely to be associated with the functions that community members have in their original host. For example, aromatic compound degradation among termite inocula is more likely to be significantly enriched because these metabolic capacities are important in fungus-growing termites, where gut microbes contribute to lignin-decomposition at the early stages of plant biomass decomposition (50), than functionally relevant when cockroaches are sustained on pure-culture fungus biomass. Further elaboration beyond *in silico* predictions will be needed to identify genes in these pathways and exploring their diet-associated expression.

Diet-imposed selective pressure on the microbiome resulted in two clearly differentiated microbiome structures (Figs. 2C; 4) and functional capabilities (Figs. 5C; 6). This flexibility is likely pivotal for the ability to track environmental changes and optimize nutrient intake or toxin degradation that maximize host fitness (3, 14, 15). This is in line with ample evidence for animals filtering their microbial community through exposure to different environments (31, 64), including via social interaction in insects (65, 66) and primates (23, 67, 68) or more intricate mechanisms such as coprophagy in wood-feeding cockroaches (24) and trophallaxis in social insects (24, 69–72). The consistent patterns support that hosts are able to plastically uptake beneficial microorganisms, thereby enhancing host fitness through its extended phenotype –its microbiota– in an environment-specific way.

The omnivorous diet imposed a stronger filter on the cockroach microbiome, which ended up being less diverse than in cockroaches fed a fungal diet. While inoculations can cause both colonizations and extinctions (i.e., colonizers can outcompete residents), nutrient uptake can solely cause extinctions (i.e., if bacteria are outcompeted on a specific set of nutrients), and cannot provoke colonizations unless microbes enter with the food. The omnivorous diet may have provoked more extinctions than the fungus diet, as it led to a less diverse microbiome (Fig. 3A). Such extinctions, occurring at a certain rate, can be considered stochastic processes which can lead the community in divergent ways, similarly to colonizations as discussed above (cf., 52). Therefore, microbiomes under the conspecific omnivorous diet, where more extinctions occur, showed less microbiome consistency than those under a specialized fungus diet (Fig. 3B). Moreover, the fact that inoculations lead to microbial colonizations while diet mostly leads to extinctions may explain the inverse correlation between diversity and microbiome divergence between diets, as opposed to between inocula and between cockroaches with or without vertically transmitted taxa, in which microbiome divergence is linked to higher diversity (Fig. 3). Therefore, diets which sustain more diverse bacteria lead to more consistent microbiota, while inocula with more diverse bacterial colonizers lead to more divergent microbiome assemblies.

Inoculation events drive communities towards different stable states depending on colonizing or extinct taxa and ecological relations between the microbiota and hosts and other gut bacteria. Conspecific inoculation opportunities offer hosts and microbes the opportunity to spend prolonged amounts of time in each-others presence, opening the door to co-evolution (2), while diet shapes the functional capabilities the microbiome houses. These processes are simultaneous, and highlight the importance of understanding microbial communities through both taxonomic and functional lenses (73–75), as ecology has done before microbiome science (76, 77). Although the relative roles of all potential factors shaping community structure will conceivably vary with hosts and diets, our findings suggest that environmental microbial sources have a stronger filtering effect on the microbiota composition than do diet, ecological relationships between early-life microbiota, or future gut colonizers. This, in turn, may lead to consistent microbial communities with host adaptations maximizing the extended phenotype that the microbiota provides (11, 78), particularly if microbes spend most of their lives in host-associated environments and, in turn, can co-evolve with hosts (2).

## Materials and Methods

### Study species

Omnivorous *S. lateralis* reproduce sexually and cover their eggs in an ootheca, which is deposited and left until hatching (Fig. 1A). Adults do not provide parental care, nymphs are self-sufficient after hatching, and obtain microbes via coprophagy and selective recruitment from the environment (31, 79, 80). After obtaining a *S. lateralis* colony from an online breeder (https://www.ebay.com/usr/zoofoods), the colony was kept in a large plastic box (57 × 40 × 20 cm) with an aluminum mesh at 27°C and 50% relative humidity (RH). Cockroaches were fed *ad libitum* with dog food (Hund, Netto A/S, Denmark) and water was supplied with soaked water-absorbing polymer crystals (http://insektorama.dk/koeb/vand-krystaller/). As an allospecific gut inoculum source, we used workers of the fungus-growing termite species *M. subhyalinus* (Termitidae, Macrotermitinae) from a laboratory colony at the Université Sorbonne Paris Nord. Termites were kept at 30°C and 75% RH, fed with dry wood and kept humid with water-soaked paper towels.

### Antimicrobial (peracetic acid) treatment of cockroaches

Twelve ootheca that produced between nine and 19 individuals were collected from the main colony. Six ootheca received an antimicrobial treatment by rinsing in 0.1% sodium dodecylbenzene sulfonate (Sigma-Aldrich, Germany, Cas number: 25155-30-0), followed by five minutes in 2% peracetic acid (Supelco, Denmark: Cas number: 79-21-0) and a rinse in 5 mL sterile water (Sigma-Aldrich, Germany, Cas number: 7732-18-5) as per the protocol by Tegtmeier et al. (41) and Mikaelyan et al. (31), who also did not find any apparent negative effects on cockroaches. Untreated ootheca were brushed free of dirt and were rinsed with 5 ml sterile water to mimic handling. Ootheca were subsequently placed individually in sterile 50 mL polypropylene Falcon tubes; cockroaches hatched from the ootheca after 27±2 days (mean ± SE; Fig. 1).

We assessed the effectiveness of the antimicrobial treatment using one cockroach per ootheca (Fig. 1; Table S1). A hatchling was crushed in 300 μL 0.2% saline solution (Sigma-Aldrich, Germany, Cas number: 7732-18-5, Sigma-Aldrich, Germany 7647-14-5), vortexed and 50 μL of the mixture was plated onto two different Potato Dextrose Agar plates (PDA; 39g/L PDA, 10 g/L agar). The remaining 200 μL was snap frozen in liquid nitrogen and stored at −20°C for MiSeq amplicon sequencing (see below). One plate was kept anoxic in a GasPak 100™ Anaerobic System (Becton-Dickinson, USA), while the other was left at ambient oxygen levels. Plates were left at room temperature for a minimum of one month and the number of Colony Forming Units (CFUs) was counted for each morphologically distinct microbe (up to 100 per unique morphology) and results were visualized in R (81) using the package ggplot (82).

### Diet manipulation and gut inoculum test

After hatching, cockroach nymphs were starved for five days within the polymer tubes and in the presence of their ootheca of origin they hatched from. Tubes were placed on ice for 15 minutes to ease handling of nymphs, prior to being individually placed into plastic boxes (7.5 cm × 5.5 cm × 5 cm) with lids that had holes covered by an aluminum mesh to allow ventilation, and a shelter made from an upside-down cardboard chicken egg-holder that was UV sterilized for 30 min prior to use. Nymphs from both antimicrobial-treated and control ootheca were distributed to one of the four treatment groups (Fig. 1). The number of nymphs used from each ootheca varied (Table S1).

Nymphs were offered a single gut inocula (cockroach or termite) in a feeding block as soon as they were moved to their experimental cages, which was removed a day later. To create feeding blocks, adult cockroaches and termite workers were placed on ice for five minutes and then euthanized by removing the head. Dissected guts (one gut per individual to be fed with the gut inoculum) were first mixed in 0.2% autoclaved saline solution and subsequently homogenized in 150 μL per gut in Potato Dextrose Broth (PDB; 39 g/L PDB) by crushing and brief vortexing. The PDB/gut mixture was then mixed with gelatin (Sigma-Aldrich, Germany, Cas number: 9000-70-8) to acquire solid feeding blocks, which were divided equally among the respective individual cages.

Respective diets were given *ad libitum* alongside the soaked water crystals. The fungal diet consisted of fresh mycelium of a *Termitomyces* species (the food source of the termite inoculum source) isolated from the fungus gardens of an *Odontotermes* cf. *badius* mound (termite colony code ICOO20) collected in Ivory Coast in 2018. The fungus was cultivated on PDA plates at room temperature until the mycelium covered the entire Petri dish, after which it was harvested. Fungus-fed cockroaches had *Termitomyces* placed onto their water crystals to avoid the fungus drying out. Individuals were fed twice per week and leftover food was removed at each feeding session. After four weeks, the cockroaches were dissected, and guts were snap-frozen in liquid nitrogen and stored at −20ÀC until DNA extraction.

### Molecular methods

DNA was extracted using the Qiagen DNeasy Blood & Tissue Kit (Qiagen, Germany), following a modified version of the manufacturer’s protocol. Frozen guts were crushed with a pestle that was subsequently rinsed with 180 μL ATL buffer. After adding glass beads, samples were vortexed for 15 s prior to adding 200 μL of chloroform:IAA. After centrifugation of samples at 20,000 g for 15 min, 80 μL of the supernatant was transferred to a sterile Eppendorf tube, and treated with 4 μL of RNase. The manufacturer’s instructions were followed during the rest of the extraction protocol. The volumes of ethanol and AL buffer were adjusted according to the digest volume (80 μL AL buffer and ethanol were added to the reaction). Finally, samples were eluted twice using 100 μL of AE buffer to maximize DNA yield.

PCR amplification of the V3-V4 region of the 16S rRNA gene using the primer set 341F-806R (5’-ACTCCTACGGGAGGCAGCAG–3’; 5’ – GGACTACHVGGGTWTCTAAT −3’) (83, 84). Initial PCRs were conducted to assess the presence of bacterial DNA. The PCR reaction mixture (10 μL) contained 1 μL template, 0.4 μl of each primer and 5 μL VWR Red Taq DNA polymerase Master mix (VWR Chemicals, Denmark, Cas nr: 733-2547). The PCR reaction conditions for amplification of DNA were as follows: initial denaturation at 94°C for 4 minutes, followed by 40 cycles of denaturation at 94°C for 30 s, annealing at 55 for 30 s and extension at 72°C for 30 s, and finally ending with 72°C for 4 min. Positive PCR was evaluated on a 2% agarose gel. Library preparation and amplicon sequencing of 123 samples were done at BGI (https://www.bgi.com/us/) and all were successful, except the blank. Samples were sequenced in two batches: the first batch included guts from the pre-experiment to evaluate antimicrobial effect, a mock community and negative controls; the second included guts from the main experiment.

### Bioinformatics and statistical analyses

Batches were handled identically using the dada2 pipeline v.1.12.1 (85) using RStudio v.3.6.1 (81). Default parameters were used aside from truncation length (275, 260), maxEE (2,3) and minOverlap set to 20. Taxonomic assignment was first performed with the manually curated Dict_db 3.0 database (30), followed by reclassification using SILVA v.1.3.2 (86) of ASVs that were not classified to the genus level. This procedure allowed 96.4% of ASVs and 99.7% of reads to be classified to genus level in batch 1 and 92.1% of ASVs and 98.4% of reads in batch 2. *Blattabacterium,* an intracellular symbiont (37), was found in all controls and these were therefore ignored. A cellular mock community standard (Zymobiomics, Nordic BioSite ApS, Copenhagen) was used to control for extraction, PCR and sequencing biases. All eight expected taxa were detected and classified appropriately and no contaminants were detected in the mock community, despite lower DNA concentrations than biological samples, strongly suggesting sample cleanliness. *Blattabacterium* and Eukaryote ASVs were filtered, as were ASVs with less than 10 observations in the full dataset. Rarefaction plots (plotted prior to rare ASV removal; Fig. S2) indicate we captured a satisfactory portion of the full bacterial community. Alpha diversity was calculated using the R package phyloseq *estimate richness* function v.1.28.0 (87). ANOVAs (88) were performed to calculate significant differences between experimental setups using the stats package v.3.6.1 (81), after testing deviation from assumptions with *shapiro.test* (89) and *bptest* (90) from the lmtest package v.0.9.37 (91). Whenever ANOVA assumptions were not met, a Kruskal Wallis was performed for univariate tests (92); or a Welch Approximate Degrees of Freedom test for multivariate tests with a Games-Howell post-hoc test were performed (93, 94). To assess community-level differences in gut microbiomes under different treatments, a Permutational Multivariate Analysis of Variance (PERMANOVA; 95) was performed using Bray-Curtis (96) distances with 10^4^ permutations using the adonis function from vegan v.2.5.6 (97). All p values were adjusted with Benjamini-Hochberg correction (98). Community level differences were visualized using non-metric multidimensional scaling (NMDS) plots using ggplot2 (82) and viridis (99) based on Bray-Curtis dissimilarity distances. Finally, we calculated within-group microbiome consistency by inferring a pergroup representative sample with the median abundance of each taxon from that group. Next, we calculated beta diversity between each sample in a group and the representative sample in that group (as in 43). Non-parametric Welch ADF tests were used to test for significant differences between diets, inocula and antimicrobial-treated vs. controls since data significantly differed from homoscedasticity.

Differentially-abundant bacterial taxa between different treatment groups were identified following CoDa good practices (44, 45, https://github.com/ggloor/CoDa_microbiome_tutorial). A center-log-ratio transformed multivariate model was conducted using the Aldex2 package (100) including variables inoculum, diet and antimicrobial treatment. After inspecting MA and volcano plots, significant values (p_adj_ < 1e-5) were extracted and plotted against their centered log ratio. As binning by genus is dependent on the order of classification database used, we repeated the analysis using SILVA first and Dict_db second, as described above (Supplementary Figure S6). Functional prediction changes associated with changing inoculum and diets were evaluated across treatment groups and on differentially abundant genera specifically using PICRUSt2 on 16S rRNA sequences (47). Subsequently, predicted microbial metabolic pathways were identified using the MetaCyc database (48).

## Data availability

MiSeq data is available from the SRA archive at NCBI (BioProject PRJNA642018). All scripts used are included as Supplementary Material.

## Acknowledgements

We thank Aram Mikaelyan, Callum Richards, and Saria Otani for fruitful discussions in the early stages of this work, Sylvia Mathiasen for help with laboratory work, Rasmus Stenbak Larsen and Pol Lannes for help with maintenance of the cockroaches, and Veronica M. Sinotte, Nick Bos, Ana Cuesta, and Blanca Ballesteros for comments on a previous draft of the manuscript. This work was supported by the European Research Council Consolidator Grant 771349 to MP. KHB was supported by a Carlsberg Foundation Distinguished Associate Professor Fellowship awarded to Assoc. Prof. Knud A. Jønsson, Natural History Museum of Denmark (CF17-0248).

## Author Contributions

KG and MP: Developed the idea and designed the experiment.

KG: Conducted experiments, laboratory work and contributed to bioinformatics analysis.

DSD: Provided termites.

JRH: Led the bioinformatics analysis.

KHB: Supervised laboratory work and contributed to bioinformatics analysis.

All authors: Contributed to writing the manuscript and approved the final version.

## Supplementary Tables and Figures

**Table S1.** Ootheca used in the experiment, hatching time, number of individuals hatching per ootheca and the number of individuals distributed across treatments in the experiment per ootheca. One cockroach per ootheca was used for the antimicrobial treatment assessment.

| Ootheca | Antimicrobial treated (Y) or controls (C) | Ootheca hatching time (days) | Individuals hatched | # of surviving / dead individuals receiving cockroach inoculum and an omnivorous diet | # of surviving / dead individuals receiving termite inoculum and an omnivorous diet | # of surviving / dead individuals receiving cockroach inoculum and fungus diet | # of surviving / dead individuals receiving termite inoculum and fungus diet |
| --- | --- | --- | --- | --- | --- | --- | --- |
| 20 | C | 37 | 9 | 2/0 | 2/0 | 0/2 | 2/0 |
| 27 | C | 19 | 11 | 2/0 | 2/0 | 1/1 | 2/0 |
| 41 | C | 28 | 11 | 2/0 | 2/0 | 1/1 | 1/1 |
| 42 | C | 28 | 16 | 3/0 | 3/0 | 2/1 | 2/1 |
| 47 | C | 32 | 13 | 3/0 | 3/0 | 1/2 | 3/0 |
| 61 | C | 22 | 17 | 3/0 | 2/1 | 1/2 | 3/0 |
| 23 | Y | 37 | 9 | 2/0 | 2/0 | 0/2 | 0/2 |
| 29 | Y | 16 | 17 | 3/0 | 3/0 | 3/0 | 2/1 |
| 30 | Y | 19 | 18 | 1/2 | 2/1 | 2/1 | 3/0 |
| 45 | Y | 34 | 9 | 2/0 | 2/0 | 1/1 | 2/0 |
| 48 | Y | 31 | 19 | 4/0 | 4/0 | 4/0 | 3/1 |
| 49 | Y | 30 | 16 | 3/0 | 3/0 | 3/0 | 3/0 |

**Table S2.** ASV absolute abundance and taxonomic classification for both batches (separate Excel file).

**Table S3.** Kruskal-Wallis tests of the effects of antimicrobial-treatment on four different alpha diversity measures.

| Response | $\chi^2$ | df | p |
| --- | --- | --- | --- |
| Observed | 2.077 | 1 | 0.1495 |
| Shannon | 4.333 | 1 | 0.0374 |
| Simpson | 4.333 | 1 | 0.0374 |
| InvSimpson | 4.333 | 1 | 0.0374 |

**Table S4.**
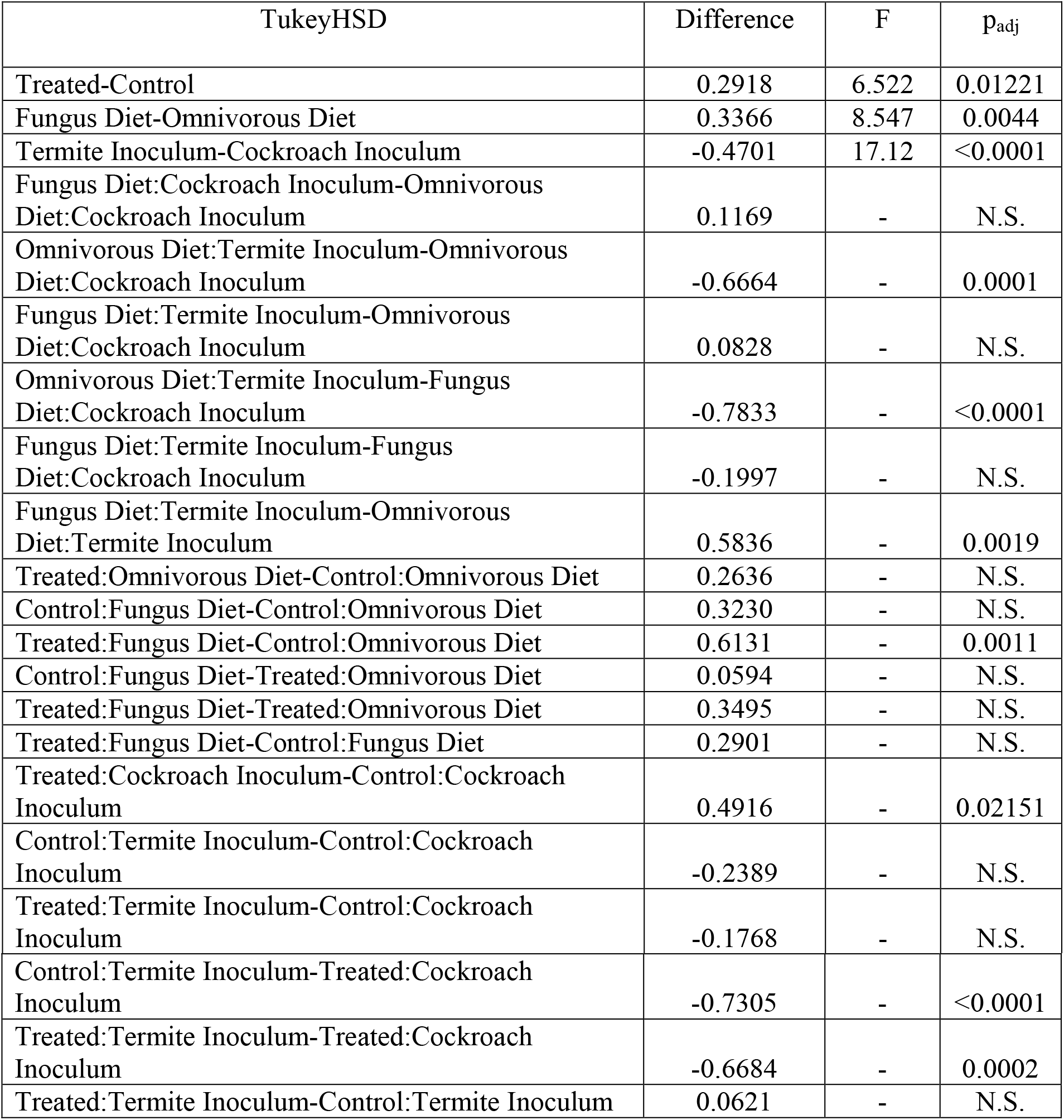
TukeyHSD post-hoc of log-transformed InvSimpson diversity multivariate ANOVA of the main experiment after removing *Blattabacterium.* F statistic from ANOVA itself.

| TukeyHSD | Difference | F | p<sub>adj</sub> |
| --- | --- | --- | --- |
| Treated-Control | 0.2918 | 6.522 | 0.01221 |
| Fungus Diet-Omnivorous Diet | 0.3366 | 8.547 | 0.0044 |
| Termite Inoculum-Cockroach Inoculum | -0.4701 | 17.12 | <0.0001 |
| Fungus Diet:Cockroach Inoculum-Omnivorous Diet:Cockroach Inoculum | 0.1169 | - | N.S. |
| Omnivorous Diet:Termite Inoculum-Omnivorous Diet:Cockroach Inoculum | -0.6664 | - | 0.0001 |
| Fungus Diet:Termite Inoculum-Omnivorous Diet:Cockroach Inoculum | 0.0828 | - | N.S. |
| Omnivorous Diet:Termite Inoculum-Fungus Diet:Cockroach Inoculum | -0.7833 | - | <0.0001 |
| Fungus Diet:Termite Inoculum-Fungus Diet:Cockroach Inoculum | -0.1997 | - | N.S. |
| Fungus Diet:Termite Inoculum-Omnivorous Diet:Termite Inoculum | 0.5836 | - | 0.0019 |
| Treated:Omnivorous Diet-Control:Omnivorous Diet | 0.2636 | - | N.S. |
| Control:Fungus Diet-Control:Omnivorous Diet | 0.3230 | - | N.S. |
| Treated:Fungus Diet-Control:Omnivorous Diet | 0.6131 | - | 0.0011 |
| Control:Fungus Diet-Treated:Omnivorous Diet | 0.0594 | - | N.S. |
| Treated:Fungus Diet-Treated:Omnivorous Diet | 0.3495 | - | N.S. |
| Treated:Fungus Diet-Control:Fungus Diet | 0.2901 | - | N.S. |
| Treated:Cockroach Inoculum-Control:Cockroach Inoculum | 0.4916 | - | 0.02151 |
| Control:Termite Inoculum-Control:Cockroach Inoculum | -0.2389 | - | N.S. |
| Treated:Termite Inoculum-Control:Cockroach Inoculum | -0.1768 | - | N.S. |
| Control:Termite Inoculum-Treated:Cockroach Inoculum | -0.7305 | - | <0.0001 |
| Treated:Termite Inoculum-Treated:Cockroach Inoculum | -0.6684 | - | 0.0002 |
| Treated:Termite Inoculum-Control:Termite Inoculum | 0.0621 | - | N.S. |

**Table S5.** Bray-Curtis multivariate PERMANOVA with 10.000 permutations after removing *Blattabacterium.*

|  | F | R<sup>2</sup> | P<sub>adj</sub> |
| --- | --- | --- | --- |
| Antimicrobial treatment | 3.422 | 0.0257 | <0.0001 |
| Diet | 8.734 | 0.0657 | <0.0001 |
| Inoculum | 15.24 | 0.1145 | <0.0001 |
| Genetic background | 1.561 | 0.1174 | <0.0001 |
| Full model |  | 0.3233 |  |

**Table S6.** Multivariate Welch Approximate Degrees of Freedom test results for intra-group microbiome divergence.

|  | WJ | df | P<sub>adj</sub> |
| --- | --- | --- | --- |
| Antimicrobial treatment | 16.81 | 1 | 0.0002 |
| Diet | 45.08 | 1 | 2.672e-8 |
| Inoculum | 8.905 | 1 | 0.0046 |
| Antimicrobial treatment:inoculum | 7.048 | 1 | 0.0109 |

**Table S7.** ALDEx2 multivariate model results table for taxa classified using DictDb first, and SILVA second (separate Excel file).

**Table S8.** ALDEx2 multivariate model results table for taxa classified using SILVA first, and DictDb second (separate Excel file).

**Table S9.** MetaCyc pathway abundance table predicted from PICRUSt2 (separate excel file).

**Table S10.** Multivariate Welch Approximate Degrees of Freedom test of predicted MetaCyc pathway from PICRUSt2 PERMANOVA analysis with 10,000 permutations.

|  | R<sup>2</sup> | F | df | P<sub>adj</sub> |
| --- | --- | --- | --- | --- |
| Antimicrobial treatment | 0.0185 | 3.081 | 1 | 0.0321 |
| Diet | 0.2224 | 37.08 | 1 | 1e-5 |
| Inoculum | 0.0618 | 10.30 | 1 | 5e-5 |
| Genetic background | 0.1578 | 2.631 | 10 | 0.0002 |

**Table S11.**
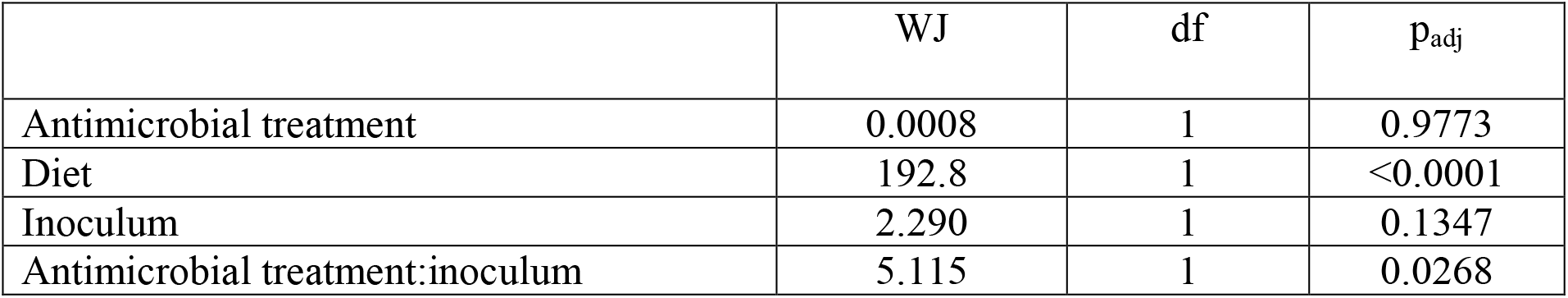
Multivariate Welch Approximate Degrees of Freedom test of predicted MetaCyc pathway from PICRUSt2 on observed richness.

**Table S12.** Multivariate Welch Approximate Degrees of Freedom test of predicted MetaCyc pathway from PICRUSt2 on InvSimpson diversity.

|  | WJ | df | P<sub>adj</sub> |
| --- | --- | --- | --- |
| Antimicrobial treatment | 2.663 | 1 | 0.1090 |
| Diet | 24.95 | 1 | 6.666e-6 |
| Inoculum | 9.468 | 1 | 0.0033 |

**Table S13.** Multivariate Welch Approximate Degrees of Freedom test of predicted MetaCyc pathway from PICRUSt2 on intra-group divergence.

|  | WJ | df | P<sub>adj</sub> |
| --- | --- | --- | --- |
| Antimicrobial treatment | 0.0759 | 1 | 0.7839 |
| Diet | 12.42 | 1 | 0.0008 |
| Inoculum | 1.284 | 1 | 0.2615 |
| Diet:inoculum | 8.251 | 1 | 0.0056 |

**Table S14.** ALDEx2 multivariate model results from predicted MetaCyc pathways from PICRUSt2 (separate Excel file).

**Table S15.**
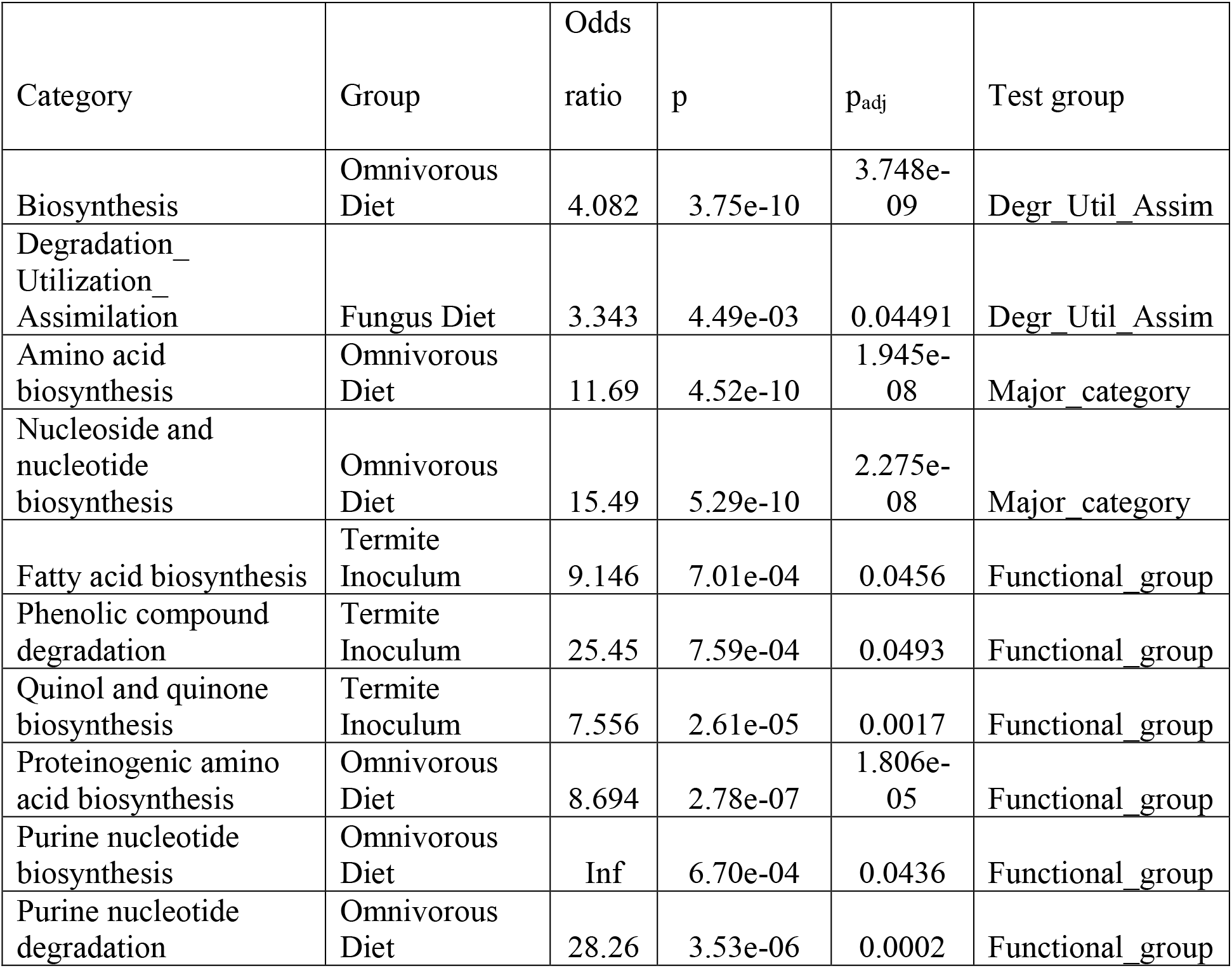
Fisher’s enrichment tests on predicted MetaCyc pathways from PICRUSt2 for all larger categories. Infinite odds ratio indicates all pathways in this group were differentially abundant in the omnivorous diet. Test group indicates the functional taxonomical level tested (MetaCyc).

**Figure S1.**
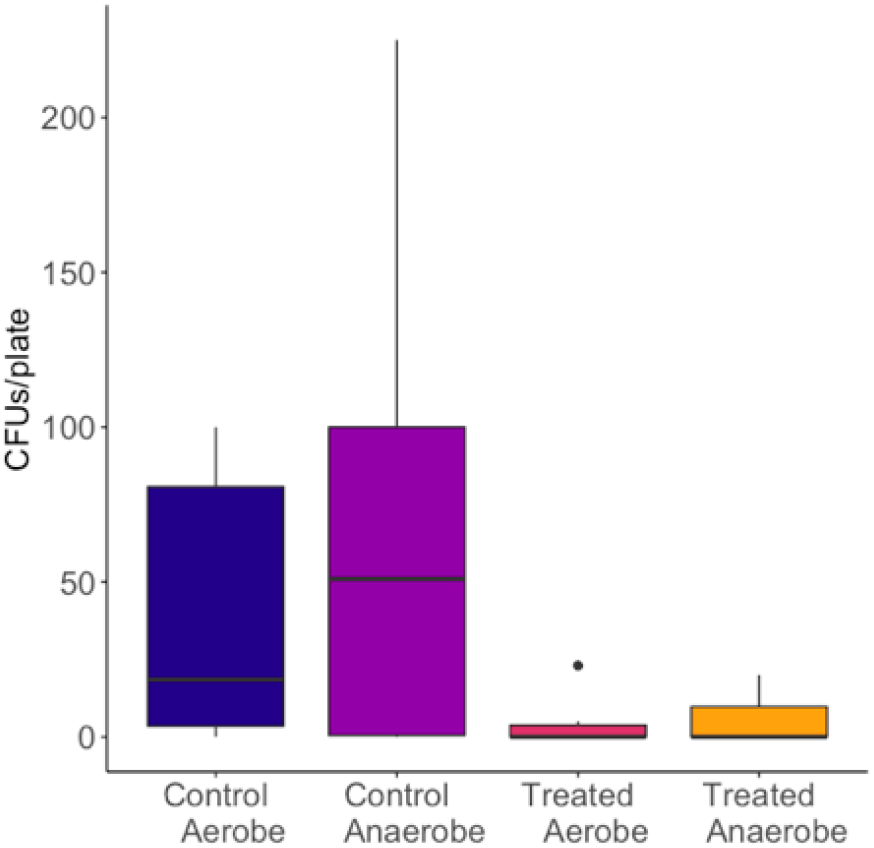
The average number of CFUs per plate from the plating experiment (n=6). Horizontal lines indicate medians, hinges indicate first and third quantiles.

**Figure S2.**
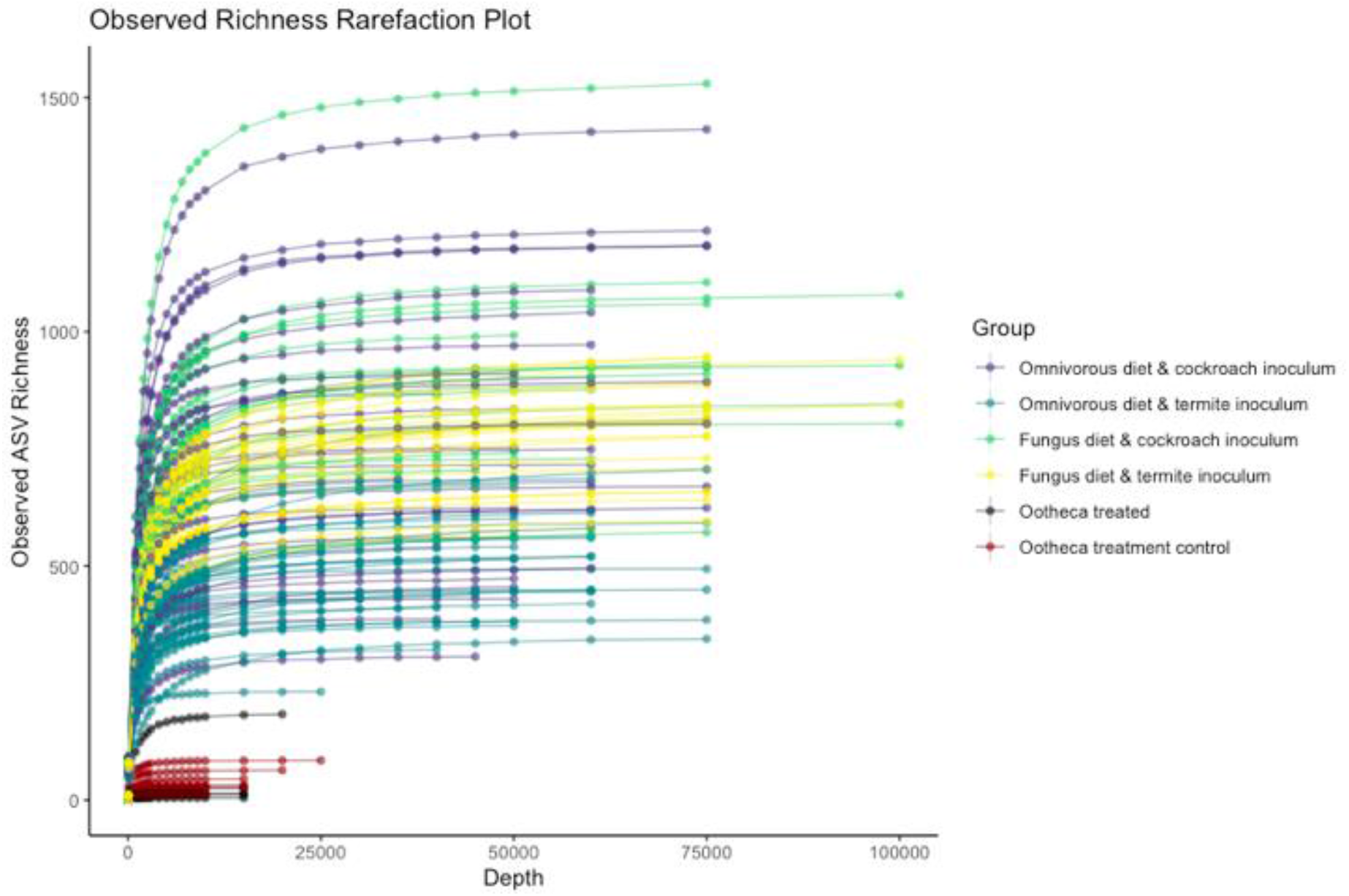
Rarefaction curves for both batches after removing *Blattabacterium*. The Y axis indicates richness at different sub-sampling sizes (X axis).

**Figure S3.**
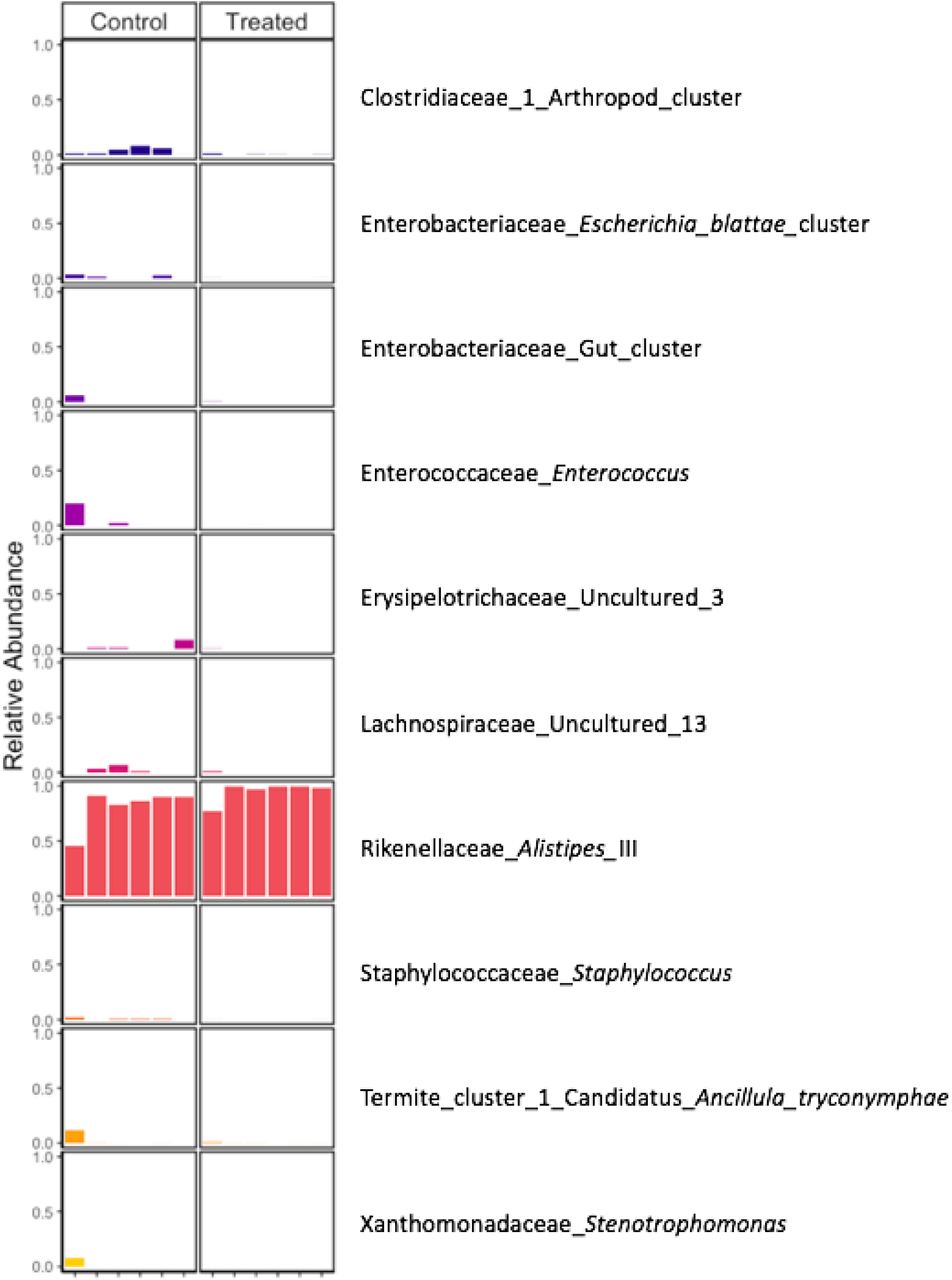
MiSeq 16S rRNA amplicon sequencing results for the test of the effectiveness of the antimicrobial treatment. The 10 most abundant genera are presented for controls (left) and antimicrobial-treated (right) microbiomes, excluding the endosymbiont *Blattabacterium*. Plot shows relative abundance for each sample.

**Figure S4.**
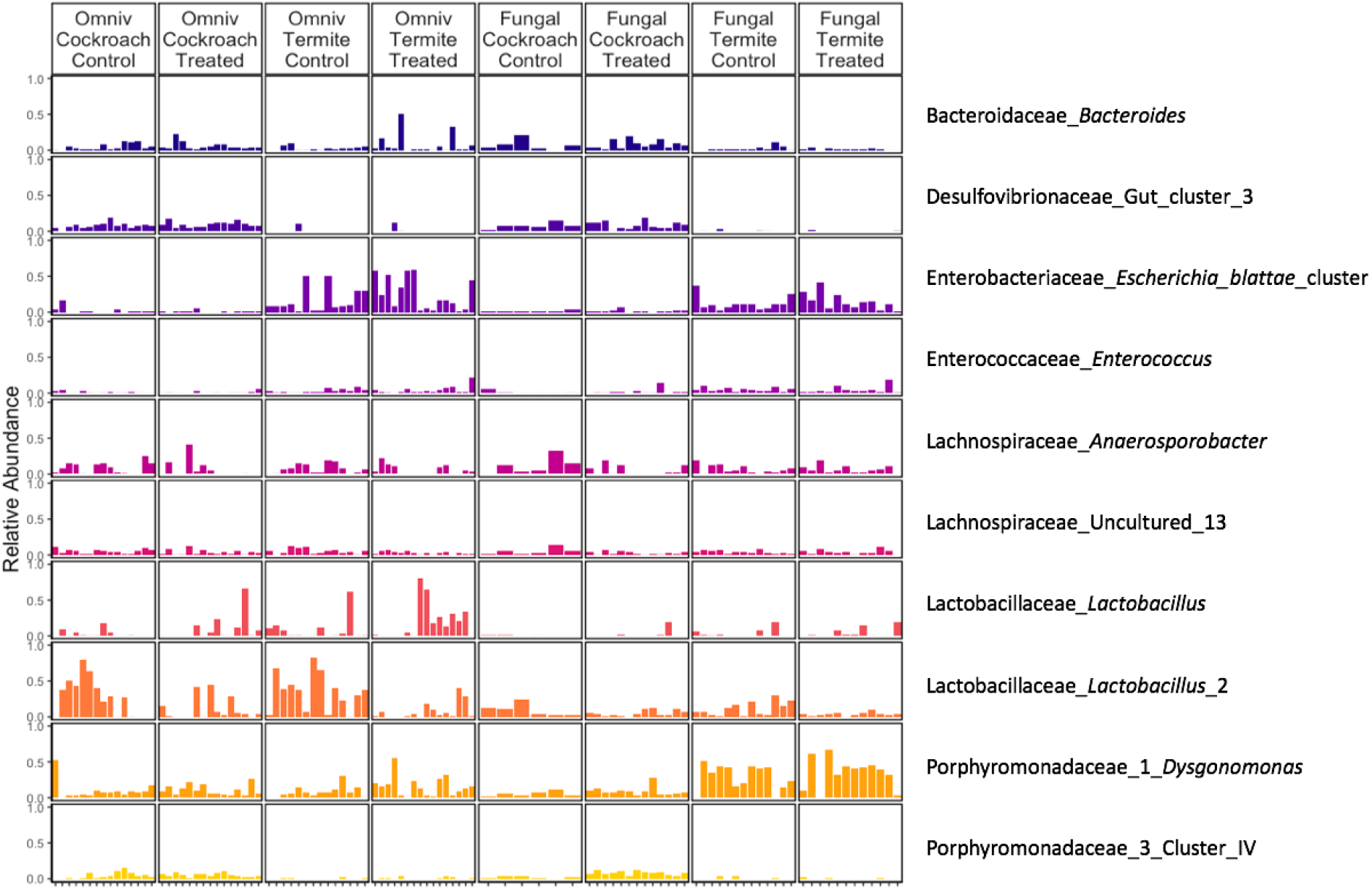
Top 10 genera across all treatment groups, sorted by relative abundance. Plot shows relative abundance for each sample. Taxon names indicate family and genus.

**Figure S5.**
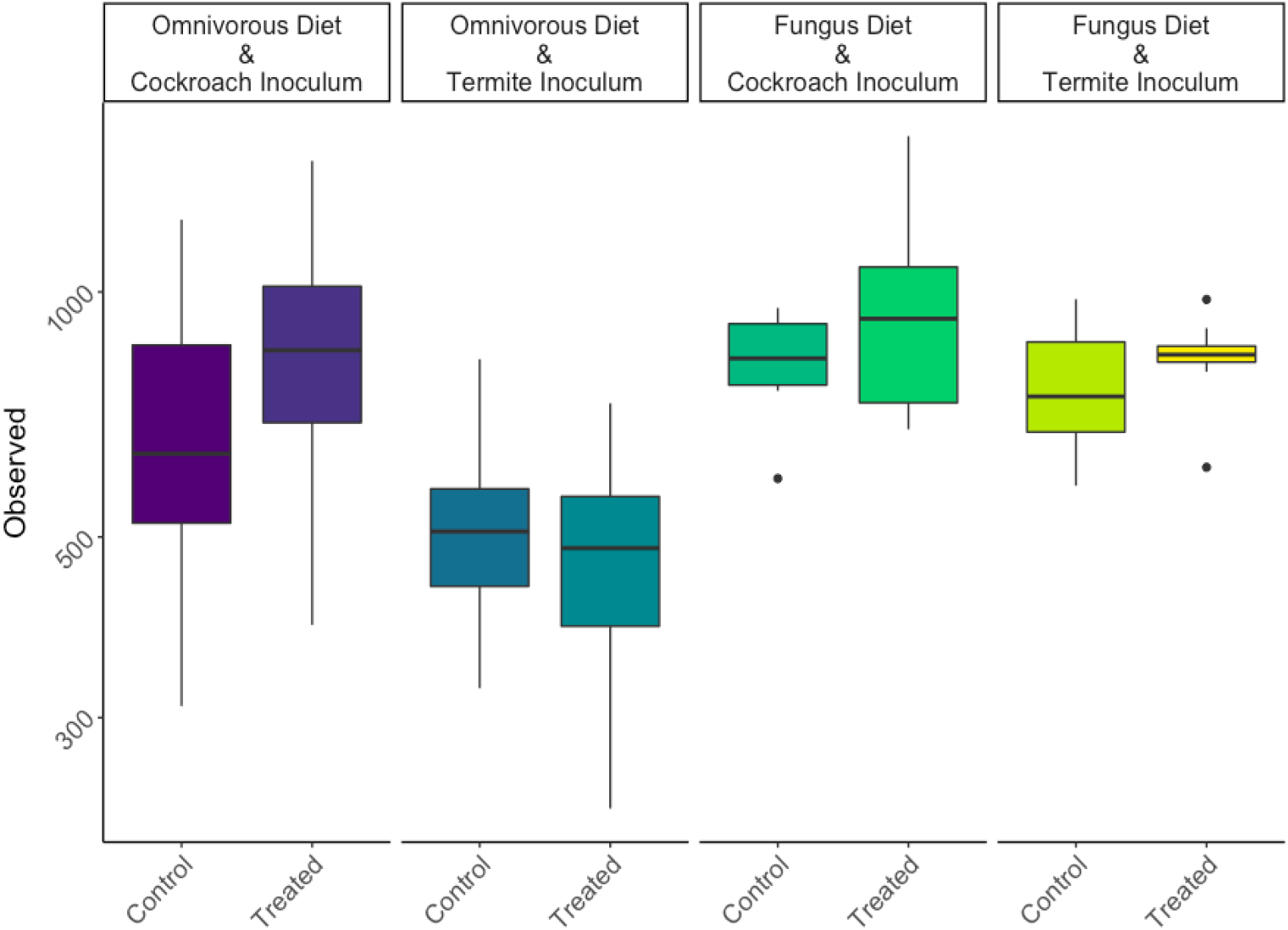
Observed richness across all eight treatment groups on a log10 scale. Horizontal lines indicate medians, hinges indicate first and third quantiles.

**Figure S6.**
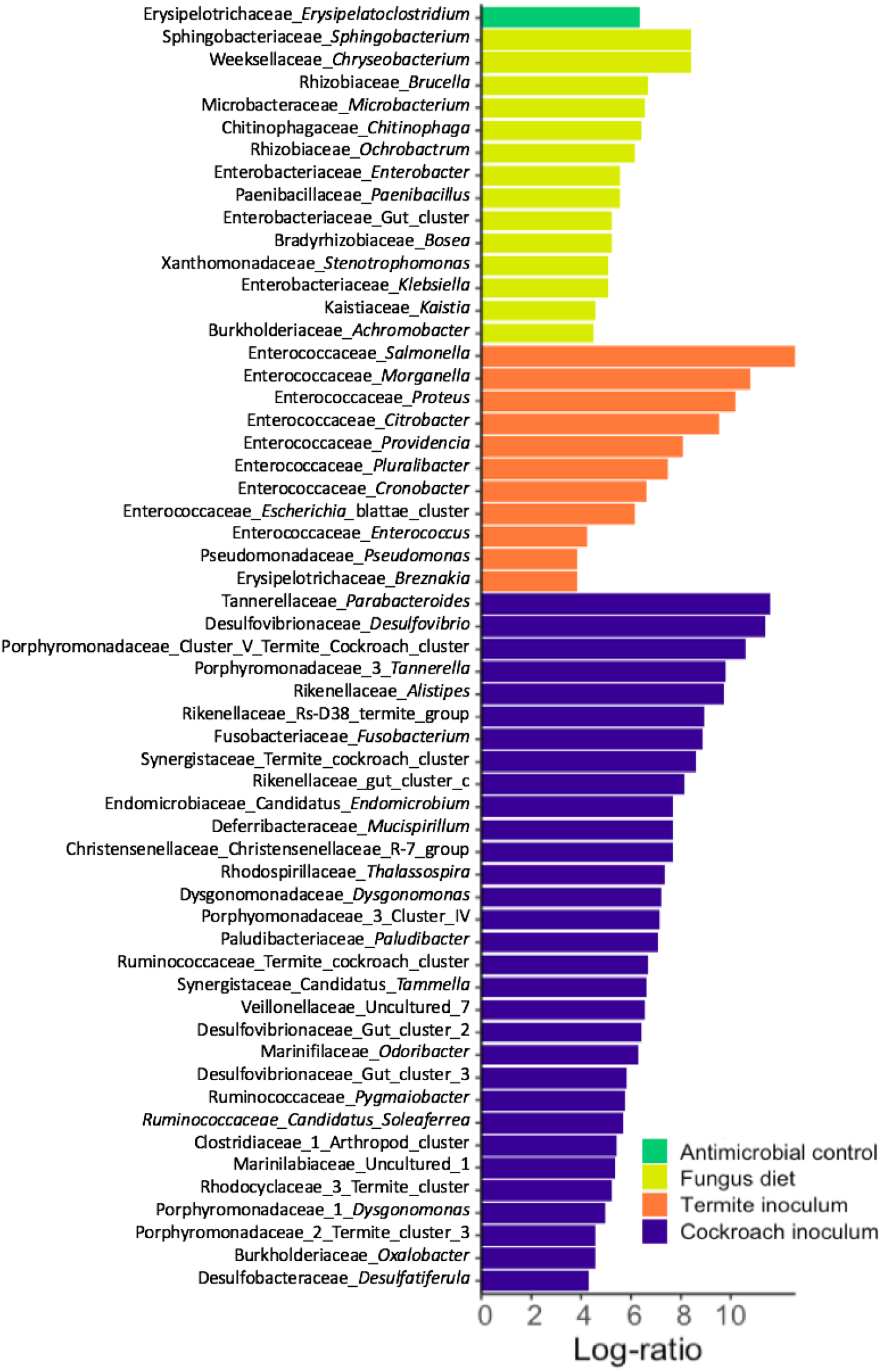
Log-ratio (effect size) increase in genera significantly correlating with inoculum, diet or antimicrobial-treated cockroaches compared to their alternative. Calculated with a multivariate Aldex2 generalized linear mixed model. All plotted taxa have Bonferroni-Hochberg adjusted p values below 1e-5. This plot differs from Fig. 4 in that classification was first done with SILVA, after which unclassified genera were reclassified with DictDb (see Materials and Methods).

**Figure S7.**
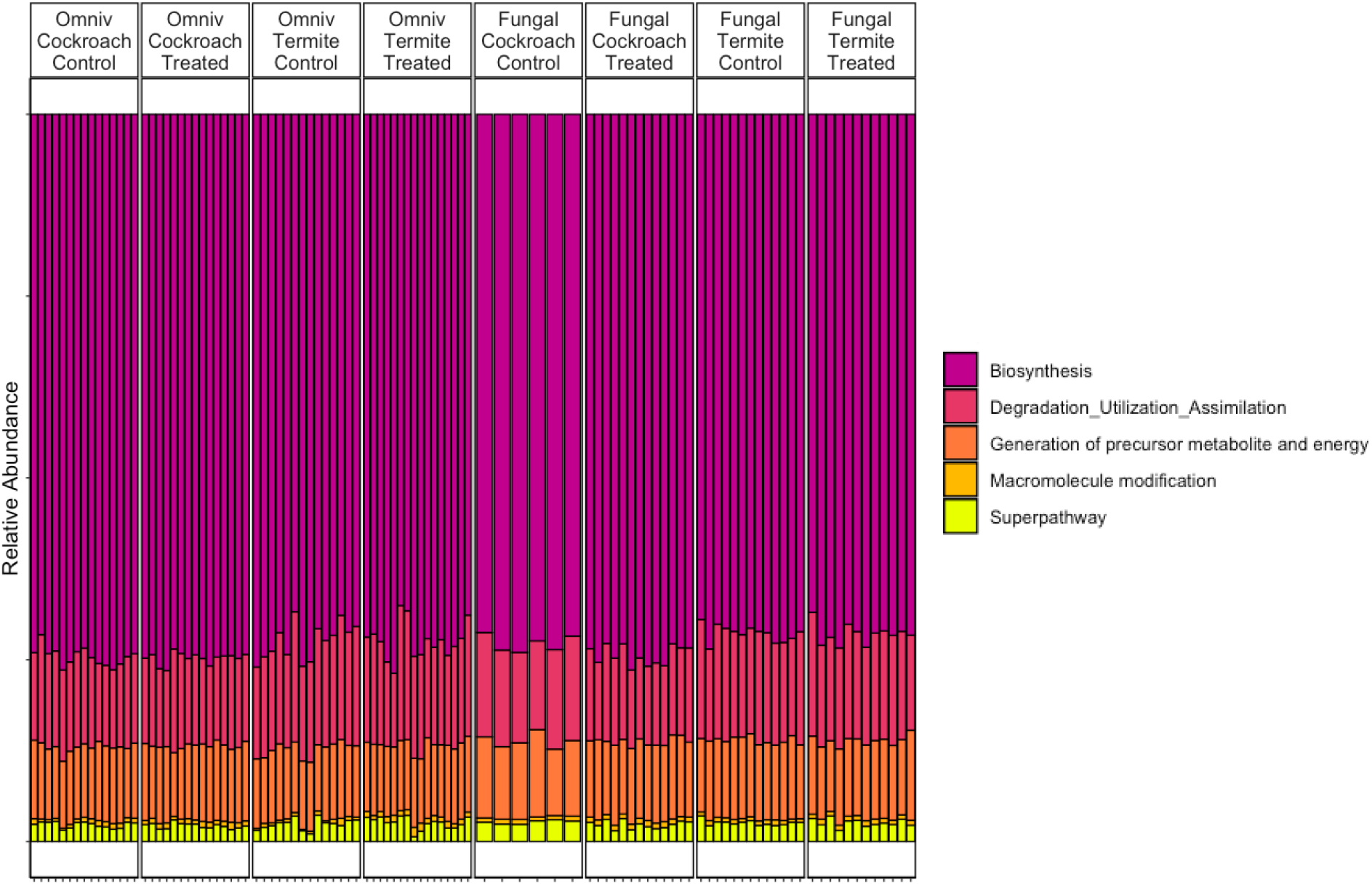
Predicted MetaCyc broadest category from PICRUSt2 pathways across treatment groups. Each column represents a sample.

**Figure S8.**
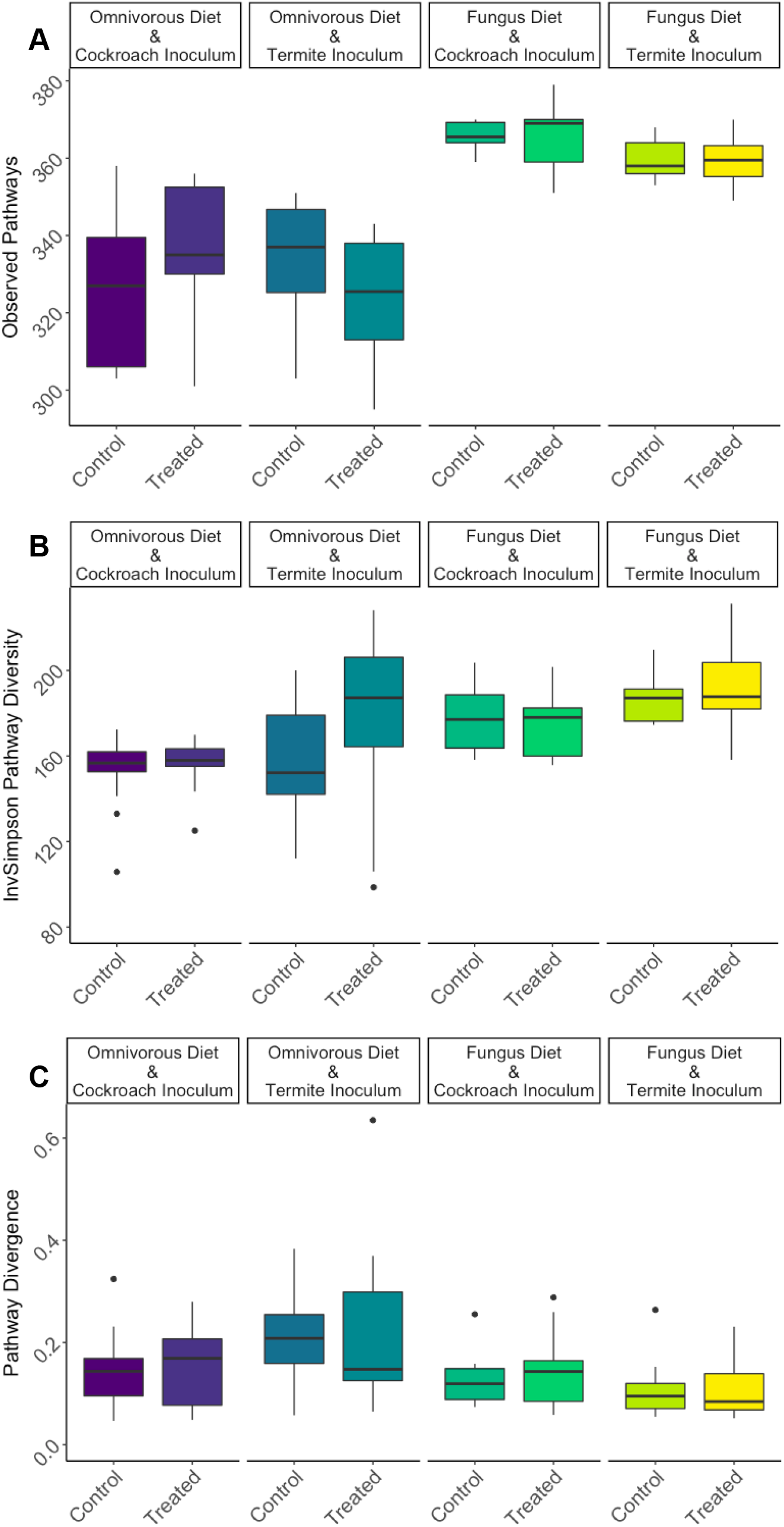
Predicted MetaCyc pathway richness (A), diversity (B) and divergence (C) of all treatment groups.

**Figure S9.**
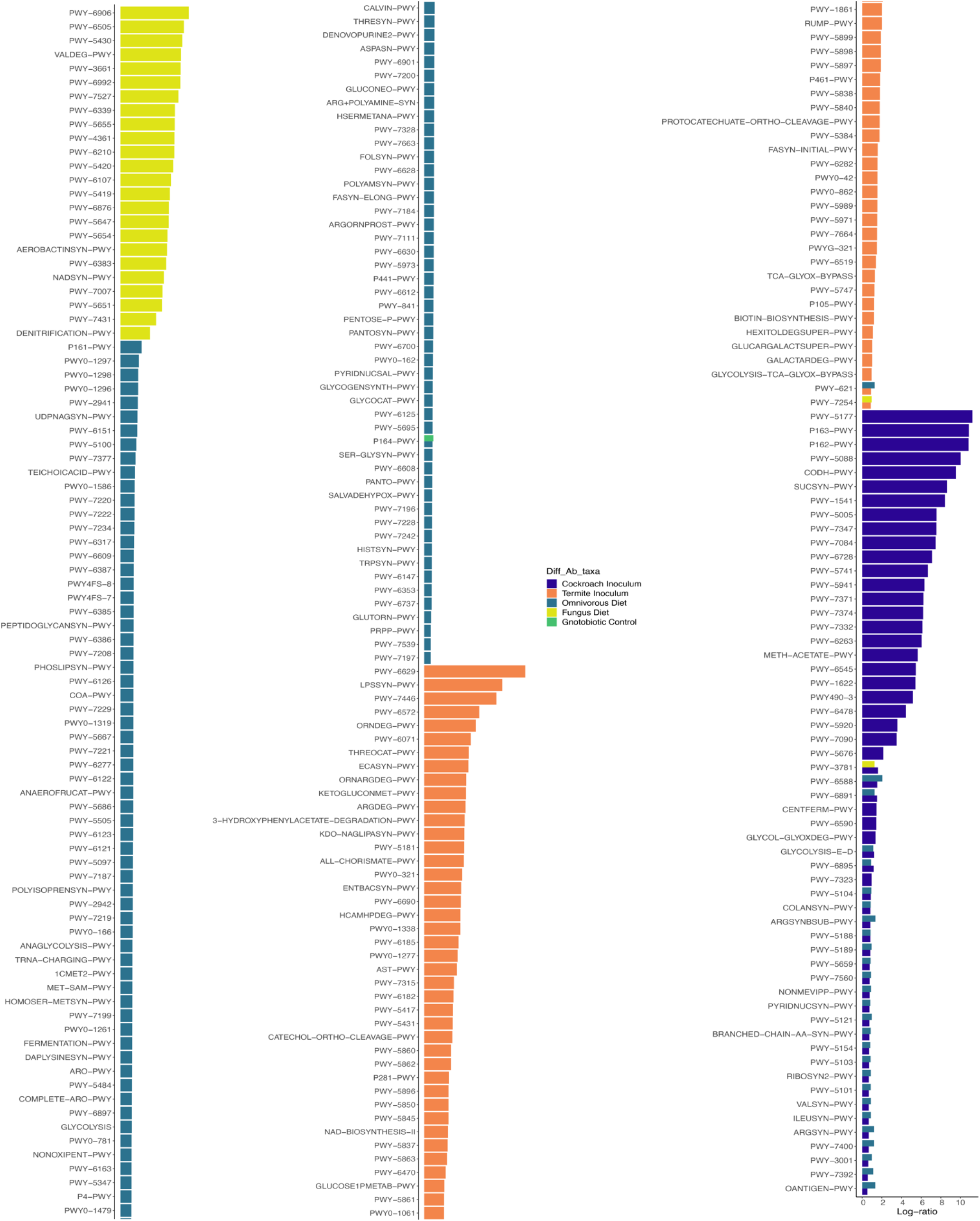
Full list of differentially abundant predicted MetaCyc pathways from PICRUSt2, between treatments. Figure split in three panels for visibility.

## References

1. Mcfall-Ngai M, Had MG, Bosch TCG, Carey H V, Domazet-lo T, Douglas AE, Dubilier N, Eberl G, Fukami T, Gilbert SF, Hentschel U, King N, Kjelleberg S, Knoll AH, Kremer N, Mazmanian SK, Metcalf JL, Nealson K, Pierce NE, Rawls JF, Reid A, Ruby EG, Rumpho M, Sanders JG, Tautz D, Wernegreen JJ. 2013. Animals in a bacterial world, a new imperative for the life sciences. Proc Natl Acad Sci U S A 110:3229–3236.

2. Moran NA, Ochman H, Hammer TJ. 2019. Evolutionary and ecological consequences of gut microbial communities. Annu Rev Ecol Evol Syst 50:451–475.

3. Wertz JT, Béchade B. 2020. Symbiont-mediated degradation of dietary carbon sources in social herbivorous insects, p. 63–109. *In* Oliver, KM, Russell, JABT-A IP (eds.), Mechanisms Underlying Microbial Symbiosis. Academic Press.

4. Michel AJ, Ward LM, Goffredi SK, Dawson KS, Baldassarre DT, Brenner A, Gotanda KM, Mccormack JE, Mullin SW, Neill AO, Tender GS, Uy JAC, Yu K, Orphan VJ, Chaves JA. 2018. The gut of the finch: uniqueness of the gut microbiome of the Galápagos vampire finch. Microbiome 6:167.

5. Moeller AH, Li Y, Mpoudi Ngole E, Ahuka-Mundeke S, Lonsdorf E V, Pusey AE, Peeters M, Hahn BH, Ochman H. 2014. Rapid changes in the gut microbiome during human evolution. Proc Natl Acad Sci U S A 111:16431–16435.

6. Delsuc F, Metcalf JL, Parfrey LW, Song SJ, González A, Knight R. 2014. Convergence of gut microbiomes in myrmecophagous mammals. Mol Ecol 23:1301–1317.

7. Anderson KE, Rusell JA, Moreau CS, Kautz S, Sullam KE, Hu Y, Basinger U, Mott BM, Buck N, Wheeler DE. 2012. Highly similar microbial communities are shared among related and trophically similar ant species. Mol Ecol 21:2282–2296.

8. Baldo L, Pretus JL, Riera JL, Musilova Z, Roger A, Nyom B, Salzburger W. 2017. Convergence of gut microbiotas in the adaptive radiations of African cichlid fishes. ISME J 11:1975–1987.

9. Dittmer J, Opstal EJ Van, Shropshire JD, Bordenstein SR, Hurst GDD, Brucker RM. 2016. Disentangling a holobiont – Recent advances and perspectives in *Nasonia* wasps. Front Microbiol 7:1478.

10. Guégan M, Zouache K, Démichel C, Minard G, Van VT, Potier P, Mavingui P, Moro CV. 2018. The mosquito holobiont: fresh insight into mosquito-microbiota interactions. Microbiome 6:49.

11. Henry LP, Bruijning M, Forsberg SKG, Ayroles JF. 2019. Can the microbiome influence host evolutionary trajectories? Biorxiv https://doi.org/10.1101/700237.

12. Sinotte VM, Renelies-Hamilton J, Taylor BA, Ellegaard KM, Sapountzis P, Vasseur-Cognet M, Poulsen M. 2020. Synergies between division of labor and gut microbiomes of social insects. Front Ecol Evol 7:503.

13. David LA, Maurice CF, Carmody RN, Gootenberg DB, Button JE, Wolfe BE, Ling A V, Devlin AS, Varma Y, Fischbach MA, Biddinger SB, Dutton RJ, Turnbaugh PJ. 2014. Diet rapidly and reproducibly alters the human gut microbiome. Nature 505:559–563.

14. Claus SP, Guillou H, Ellero-Simatos S. 2016. The gut microbiota: a major player in the toxicity of environmental pollutants? Nat Publ Gr 4: 16003.

15. Feng P, Ye Z, Han H, Ling Z, Zhou T, Zhao S, Virk AK, Kakade A, Abomohra AE, El-Dalatony MM, Salama E, Liu P, Li X. 2020. Tibet plateau probiotic mitigates chromate toxicity in mice by alleviating oxidative stress in gut microbiota. Commun Biol 3:242.

16. Macke E, Tasiemski A, Massol F, Callens M, Decaestecker E. 2017. Life history and eco-evolutionary dynamics in light of the gut microbiota. Oikos 126:508–531.

17. Dietrich C, Köhler T, Brune A, Dietrich C, Köhler T, Brune A. 2014. The cockroach origin of the termite gut microbiota: patterns in bacterial community structure reflect major evolutionary events. Appl Environ Microbiol 80: 2261–2269.

18. Groussin M, Mazel F, Sanders JG, Smillie CS, Thuiller W, Alm EJ. 2017. Unraveling the processes shaping mammalian gut microbiomes over evolutionary time. Nat Commun 8:14319.

19. Lutz HL, Jackson EW, Webala PW, Babyesiza WS, Peterhans CK, Demos TC, Patterson BD, Gilbert AJ. 2019. Ecology and host identity outweigh evolutionary history in shaping the bat microbiome. mSystems 4:e00511–e00519.

20. Jin Song S, Sanders JG, Delsuc F, Metcalf J, Amato K, Taylor MW, Mazel F, Lutz HL, Winker K, Graves GR, Humphrey G, Gilbert JA, Hackett SJ, White KP, Skeen HR, Kurtis SM, Withrow J, Braile T, Miller M, Mccracken KG, Maley JM, Ezenwa VO, Williams A, Blanton JM, McKenzie VJ, Knight R. 2020. Comparative analyses of vertebrate gut microbiomes reveal convergence between birds and bats. mBio 11: e02901–19.

21. Teyssier A, Lens L, Matthysen E, White J. 2018. Dynamics of gut microbiota diversity during the early development of an avian host: evidence from a cross-foster experiment. Front Microbiol 9:1524.

22. Lewis WB, Moore FR, Wang S. 2016. Characterization of the gut microbiota of migratory passerines during stopover along the northern coast of the Gulf of Mexico. J Avian Biol 47: 659–668.

23. Moeller AHA, Foerster S, Wilson ML, Pusey AE, Hahn BH, Ochman H. 2016. Social behavior shapes the chimpanzee pan-microbiome. Sci Adv 2: e1500997.

24. Nalepa CA, Bignell DE, Bandi C. 2001. Detritivory, coprophagy, and the evolution of digestive mutualisms in Dictyoptera. Insectes Soc 48:194–201.

25. Kobayashi A, Tsuchida S, Ueda A, Yamada T, Murata K, Nakamura H, Ushida K. 2019. Role of coprophagy in the cecal microbiome development of an herbivorous bird Japanese rock ptarmigan. J Vet Med Sci 81:1389–1399.

26. Nalepa CA. 1990. Early development of nymphs and establishment of hindgut symbiosis in *Cryptocercus punctulatus* (Dictyoptera: Cryptocercidae). Entomol Soc Am 83:786–789.

27. Fukami T. 2015. Historical contingency in community assembly: integrating niches, species pools, and priority effects. Annu Rev Ecol Evol Syst 46:1–23.

28. Sprockett D, Fukami T, Relman DA. 2018. Role of priority effects in the early-life assembly of the gut microbiota. Nat Rev Gastroenterol Hepatol 15:197–205.

29. Martínez I, Maldonado-Gomez MX, Gomes-Neto JC, Kittana H, Ding H, Schmaltz R, Joglekar P, Cardona RJ, Marsteller NL, Kembel SW, Benson AK, Peterson DA, Ramer-tait AE, Walter J. 2018. Experimental evaluation of the importance of colonization history in early-life gut microbiota assembly. Elife 7: e36521.

30. Mikaelyan A, Köhler T, Lampert N, Rohland J, Boga H, Meuser K, Brune A. 2015. Classifying the bacterial gut microbiota of termites and cockroaches: A curated phylogenetic reference database (DictDb). Syst Appl Microbiol 38:472–482.

31. Mikaelyan A, Thompson CL, Hofer MJ, Brune A. 2016. The deterministic assembly of complex bacterial communities in germ-free cockroach guts. Appl Environ Microbiol 82:1256–1263.

32. Mazel F, Davis KM, Loudon A, Kwong WK, Groussin M, Parfrey LW. 2018. Is host filtering the main driver of phylosymbiosis across the tree of life? mSystems 3: e00097–18.

33. Loo WT, García-Loor J, Rachael YD, Kleindorfel S, Cavanaugh CM. 2019. Host phylogeny, diet, and habitat differentiate the gut microbiomes of Darwin’s finches on Santa Cruz Island. Sci Rep 9:18781.

34. Lampert N, Mikaelyan A, Brune A. 2019. Diet is not the primary driver of bacterial community structure in the gut of litter-feeding cockroaches. BMC Microbiol 19:238.

35. Pérez-Cobas AE, Maiques E, Angelova A, Carrasco P, Moya A, Latorre A. 2015. Diet shapes the gut microbiota of the omnivorous cockroach *Blattella germanica*. FEMS Microbiol Ecol 91: fiv022.

36. Richards C, Otani S, Mikaelyan A, Poulsen M. 2017. *Pycnoscelus surinamensis* cockroach gut microbiota respond consistently to a fungal diet without mirroring those of fungus-farming termites. PLoS One 12: e0185745.

37. Gontang EA, Aylward FO, Carlos C, Glavina T, Chovatia M, Fern A, Lo C, Malfatti SA, Tringe G, Currie CR, Kolter R. 2017. Major changes in microbial diversity and community composition across gut sections of a juvenile *Panchlora* cockroach. PLoS One 12: e0177189.

38. Rosas T, García-Ferris C, Domínguez-Santos R, Llop P, Latorre A, Moya A. 2018. Rifampicin treatment of *Blattella germanica* evidences a fecal transmission route of their gut microbiota. FEMS Microbiol Ecol 94: fiy002.

39. Jahnes BC, Herrmann M, Sabree ZL. 2019. Conspecific coprophagy stimulates normal development in a germ-free model invertebrate. PeerJ 7: e6914.

40. Otani S, Mikaelyan A, Nobre T, Hansen LH, Koné NA, Sørensen SJ, Aanen DK, Boomsma JJ, Brune A, Poulsen M. 2014. Identifying the core microbial community in the gut of fungus-growing termites. Mol Ecol 23:4631–4644.

41. Tegtmeier D, Thompson CL, Schauer C, Brune A. 2015. Oxygen affects colonization and metabolic activities of gut bacteria in a gnotobiotic cockroach model. Appl Environ Microbiol 82:1080–1089.

42. Domínguez-Santos R, Pérez-cobas AE, Artacho A, Castro JA, Talón I, Moya A, García-ferris C, Latorre A. 2020. Unraveling assemblage, functions and stability of the gut microbiota of *Blattella germanica* by antibiotic treatment. Front Microbiol 11:487.

43. Salonen A, Lahti L, Saloja J, Holtrop G, Korpela K, Duncan SH, Date P, Farquharson F, Johnstone AM, Lobley GE, Louis P, Flint HJ, Vos WM De. 2014. Impact of diet and individual variation on intestinal microbiota composition and fermentation products in obese men. ISME J 8:2218–2230.

44. Gloor GB, Macklaim JM, Fernandes AD. 2016. Displaying variation in large datasets: plotting a visual summary of effect sizes. J Comput Graph Stat 25:971–979.

45. Gloor GB, Macklaim JM, Pawlowsky-glahn V, Egozcue JJ. 2017. Microbiome datasets are compositional: and this is not optional. Front Microbiol 8:2224.

46. Bourguignon T, Lo N, Dietrich C, Šobotník J, Sidek S, Roisin Y, Brune A, Evans TA. 2018. Rampant host switching shaped the termite gut. Curr Biol 28:649–654.

47. Douglas GM, Maffei VJ, Zaneveld J, Yurgel SN, Brown JR, Taylor CM, Huttenhower C, Langille MGI. 2019. PICRUSt2: An improved and extensible approach for metagenome inference. BioRxiv 1–42 https://doi.org/10.1101/672295.

48. Caspi R, Foerster H, Fulcher CA, Hopkinson R, Ingraham J, Kaipa P, Krummenacker M, Paley S, Pick J, Rhee SY, Tissier C, Zhang P, Karp PD. 2006. MetaCyc: a multiorganism database of metabolic pathways and enzymes. Nucleic Acids Res 34:511–516.

49. Lozupone CA, Stombaugh JI, Gordon JI, Jansson JK, Knight R. 2012. Diversity, stability and resilience of the human gut microbiota. Nature 489:220–230.

50. Li H, Yelle DJ, Li C, Yang M, Ke J, Zhang R, Liu Y, Zhu N, Liang S, Mo X, Ralph Jo, Currie CR, Mo J. 2017. Lignocellulose pretreatment in a fungus-cultivating termite. Proc Natl Acad Sci U S A 114:4709–4714.

51. Houghteling PD, Walker WA. 2016. Why is initial bacterial colonization of the intestine important to the infant’s and child’s health? J Pediatr Gastroenterol Nutr 60:294–307.

52. Obadia B, Güvener ZT, Zhang V, Ceja-navarro JA, Eoin L, Ja WW, Ludington WB. 2018. Probabilistic invasion underlies natural gut microbiome stability. Curr Biol 27:1999–2006.

53. Coyte KZ, Schluter J, Foster KR. 2015. The ecology of the microbiome: Networks, competition, and stability. Science 350:663–666.

54. Bauer MA, Kainz K, Carmona-Gutierrez D, Madeo F. 2018. Microbial wars: Competition in ecological niches and within the microbiome. Microb Cell 5:215–219.

55. Troyer K. 1984. Microbes, herbivory and the evolution social behavior. J Theor Biol 106:157–169.

56. Kopanic Jr RJ, Holbrook GL, Sevala V, Schal C. 2001. An adaptive benefit of facultative coprophagy in the German cockroach *Blattella germanica*. Ecol Entomol 26:154–162.

57. Lombardo MP. 2008. Access to mutualistic endosymbiotic microbes: an underappreciated benefit of group living. Ecol Sociobiol 62:479–497.

58. Nalepa CA. 2015. Origin of termite eusociality: trophallaxis integrates the social, nutritional, and microbial environments. Ecol Entomol 40:323–335.

59. Tinker KA, Ottesen EA. 2020. Phylosymbiosis across deeply diverging lineages of omnivorous cockroaches (order Blattodea). Appl Environ Microbiol 86: e02513–19.

60. Imhoff JF. 2005. Enterobacteriales, p. 587–850. *In* Brenner (ed.), Bergey’s Manual of Systematic Bacteriology. Springer Boston.

61. Tian J, Pourcher A, Peu P. 2016. Isolation of bacterial strains able to metabolize lignin and lignin-related compounds. Lett Appl Microbiol 63:30–37.

62. Taylor BF. 1983. Aerobic and anaerobic catabolism of vanillic acid and some other methoxy-aromatic compounds by *Pseudomonas* sp. strain PN-1. Appl Environ Microbiol 46:1286–1292.

63. Liang L, Song X, Kong J, Shen C, Huang T, Hu Z. 2014. Anaerobic biodegradation of high-molecular-weight polycyclic aromatic hydrocarbons by a facultative anaerobe *Pseudomonas* sp. JP1. Biodegradation 25:825–833.

64. Moeller AH, Peeters M, Ndjango JB, Li Y, Hahn BH, Ochman H. 2013. Sympatric chimpanzees and gorillas harbor convergent gut microbial communities. Genome Res 23:1715–1720.

65. Engel P, Moran NA. 2013. The gut microbiota of insects – diversity in structure and function. FEMS Microbiol Rev 37:699–735.

66. Kwong WK, Engel P, Koch H, Moran NA. 2014. Genomics and host specialization of honey bee and bumble bee gut symbionts. Proc Natl Acad Sci U S A. 111:11509–11514.

67. Tung J, Barreiro LB, Burns MB, Grenier JC, Lynch J, Grieneisen LE, Altmann J, Alberts SC, Blekhman R, Archie EA. 2015. Social networks predict gut microbiome composition in wild baboons. Elife 2015: e05224.

68. Bennett G, Malone M, Sauther ML, Cuozzo FP, White B, Nelson KE, Stumpf RM, Knight R, Leigh SR, Amato KR. 2016. Host age, social group, and habitat type influence the gut microbiota of wild ring-tailed lemurs *(Lemur catta)*. Am J Primatol 78:883–892.

69. Wheeler DE. 1984. Behavior of the ant, *Procryptocerus scabriusculus* (Hymenoptera: Formicidae), with comparisons to other Cephalotines. Psyche (Stuttg) 91:171–192.

70. Ohkuma M., Brune A. 2010. Diversity, Structure, and Evolution of the Termite Gut Microbial Community, p. 431–438. *In:* Bignell D., Roisin Y., Lo N. (eds) Biology of Termites: A Modern Synthesis. Springer, Dordrecht

71. Powell JE, Martinson VG, Urban-mead K, Moran A. 2014. Routes of acquisition of the gut microbiota of the honey bee *Apis mellifera*. Appl Environ Microbiol 80:7378–7387.

72. Lanan MC, Augusto P, Rodrigues P, Agellon A, Jansma P, Wheeler DE. 2016. A bacterial filter protects and structures the gut microbiome of an insect. Nat Publ Gr 10:1866–1876.

73. Blondel J. 2003. Guilds or functional groups: does it matter? OIKOS 2:223–231.

74. Inkpen SA, Douglas GM, Brunet TDP, Leuschen K, Doolittle WF, Langille MGI. 2017. The coupling of taxonomy and function in microbiomes. Biol Philos 32:1225–1243.

75. Banerjee S, Schlaeppi K, Heijden MGA. 2018. Keystone taxa as drivers of microbiome structure and functioning. Nat Rev Microbiol 16:567–576.

76. Simberloff D. 1991. The guild concept and the structure of ecological communities. Annu Rev Ecol Evol Syst 22:115–143.

77. Prosser JI, Bohannan BJM, Curtis TP, Ellis RJ, Firestone MK, Freckleton RP, Green JL, Green LE, Killham K, Lennon JJ, Osborn AM, Solan M, Gast CJ Van Der, Young JPW. 2007. The role of ecological theory in microbial ecology. Nature 5:384–392.

78. Foster KR, Schluter J, Coyte KZ, Rakoff-Nahoum S. 2017. The evolution of the host microbiome as an ecosystem on a leash. Nature 548:43–51.

79. Schauer C, Thompson CL, Brune A. 2012. The bacterial community in the gut of the cockroach *Shelfordella lateralis* reflects the close evolutionary relatedness of cockroaches. Appl Environ Microbiol 78:2758–2767.

80. Schauer C, Thompson C, Brune A. 2014. Pyrotag sequencing of the gut microbiota of the cockroach *Shelfordella lateralis* reveals a highly dynamic core but only limited effects of diet on community structure. PLoS One 9: e85861.

81. R Core Team. 2019. R: A language and environment for statistical computing.

82. Wickham H. 2016. ggplot2: elegant graphics for data analysis.

83. Caporaso JG, Lauber CL, Walters WA, Berg-Lyons D, Lozupone CA, Turnbaugh PJ, Fierer N, Knight R. 2010. Global patterns of 16S rRNA diversity at a depth of millions of sequences per sample. Proc Natl Acad Sci U S A 108:4516–4522.

84. Takahashi S, Tomita J, Nishioka K, Hisada T, Nishijima M. 2014. Development of a prokaryotic universal primer for simultaneous analysis of bacteria and archaea using nextgeneration sequencing. PLoS One 9: e105592.

85. Callahan BJ, McMurdie PJ, Rosen MJ, Han AW, Johnson, AJA, Holmes SP. 2016. DADA2: High-resolution sample inference from Illumina amplicon data. Nat Meth 13:581–583.

86. Quast C, Pruesse E, Yilmaz P, Gerken J, Schweer T, Yarza P, Peplies J, Glöckner O. 2013. The SILVA ribosomal RNA gene database project: improved data processing and webbased tools. Nucleic Acids Res 41:590–596.

87. McMurdie PJ, Holmes S. 2013. Phyloseq: A bioconductor package for handling and analysis of high-throughput phylogenetic sequence data. Pac Symp Biocomput 235–246.

88. Chambers J M, Freeny A, Heiberger R M. 1992. Analysis of variance; designed experiments, p 145–193. *In* Statistical Models in S. Chambers JM and Hastle TJ (eds), Wadsworth & Brooks/Cole.

89. Royston P. 1982. An extension of Shapiro and Wilk’s W test for normality to large samples. Appl Stat 31:115–124.

90. Breusch T S, Pagan A R. 1979. A simple test for heteroscedasticity and random coefficient variation. Econometrica 57:1287–1294

91. Zeileis A, Hothorn T. 2002. Diagnostic checking in regression relationships. R News. 2:7–10.

92. Hollander M, Wolfe D A. 1973. Nonparametric statistical methods. John Wiley & Sons. 115–120.

93. Efron B, Tibshirani R. 1986. Bootstrap methods for standard errors, confidence intervals and other measures of statistical accuracy. Stat Sci, 1:54–75.

94. Villacorta P J. 2017. The welchADF Package for robust hypothesis testing in unbalanced multivariate mixed models with heteroscedastic and non-normal data. The R Journal, 9:309–328.

95. Legendre P, Anderson M J. 1999. Distance-based redundancy analysis: testing multispecies responses in multifactorial ecological experiments. Ecol Monogr 69:1–24.

96. Bray JR, Curtis JT. 1957. An ordination of the upland forest communities of Southern Wisconsin. Ecol Monogr 27:325–349.

97. Oksanen J, Blanchet F G, Friendly M, Kindt R, Legendre P, McGlinn D, Minchin P R, O’Hara R B, Simpson G L, Solymos P, Stevens M H H, Szoecs E, Wagner H. 2018. Vegan: community ecology package.

98. Benjamini Y, Hochberg Y. 1995. Controlling the false discovery rate: a practical and powerful approach to multiple testing. J R Stat Soc 57:289–300.

99. Garnier S, Ross, N, Rudis, B, Sciaini, M, Scherer, C. 2018. Viridis: default color maps from matplotlib. https://cran.r-project.org/web/packages/viridis/index.html

100. Fernandes A D, Macklaim J M, Linn T G, Reid G, Gloor G B. 2013. ANOVA-like differential gene expression. analysis of single-organism and meta-RNA-seq. PLOS ONE. 8:e67019.

